# mosGraphFlow: a novel integrative graph AI model mining disease targets from multi-omic data

**DOI:** 10.1101/2024.08.01.606219

**Authors:** Heming Zhang, Dekang Cao, Tim Xu, Emily Chen, Guangfu Li, Yixin Chen, Philip Payne, Michael Province, Fuhai Li

## Abstract

Multi-omic data can better characterize complex cellular signaling pathways from multiple views compared to individual omic data. However, integrative multi-omic data analysis to rank key disease biomarkers and infer core signaling pathways remains an open problem. In this study, our novel contributions are that we developed a novel graph AI model, *mosGraphFlow*, for analyzing multi-omic signaling graphs (mosGraphs), 2) analyzed multi-omic mosGraph datasets of AD, and 3) identified, visualized and evaluated a set of AD associated signaling biomarkers and network. The comparison results show that the proposed model not only achieves the best classification accuracy but also identifies important AD disease biomarkers and signaling interactions. Moreover, the signaling sources are highlighted at specific omic levels to facilitate the understanding of the pathogenesis of AD. The proposed model can also be applied and expanded for other studies using multi-omic data. Model code is accessible via GitHub: https://github.com/FuhaiLiAiLab/mosGraphFlow

## Introduction

The advent of multi-omic data has revolutionized the field of biomedical research by providing a comprehensive view of the complex biological processes underlying various diseases. Unlike single-omic approaches, which focus on a specific type of molecular data such as genomics, transcriptomics, or proteomics, multi-omic data integrates information from multiple molecular layers to measure and characterize the multi-level molecular genotype of diseases. This integrative approach offers a more holistic understanding of cellular signaling pathways, enabling researchers to uncover intricate molecular interactions and regulatory mechanisms. Multi-omic datasets have proven invaluable in identifying essential disease biomarkers and elucidating dysfunctional signaling pathways, particularly in understanding the genetic heterogeneity of diseases at multiple levels. Despite its potential and demonstrated utility, the effective integration and analysis of multi-omic data to identify key disease biomarkers and elucidate core signaling pathways remain significant challenges. Traditional AI models often struggle to fully leverage the richness of multi-omic data due to its complexity and high dimensionality. However, recent advancements in graph AI models have shown promise in addressing these challenges by utilizing graph-based representations to capture the intricate relationships within multi-omic datasets, offering new avenues for biomarker discovery and pathway inference. This approach can be instrumental in enhancing our understanding of disease pathogenesis and in designing more effective therapeutic interventions.

Alzheimer’s disease (AD) is the most prevalent cause of dementia, primarily affecting individuals over the age of 65, though cases in younger individuals starting from around age 40 are increasingly observed. Characterized by progressive cognitive impairment, AD manifests through the hallmark neuropathological features, extracellular amyloid-β plaques and intracellular neurofibrillary tangles (NFT), caused by amyloid-β accumulation and tau hyperphosphorylation^1^. Linked to these hallmarks are blood-brain barrier disruption, mitochondrial impairment, neuroinflammation, synaptic impairment and neuronal loss. The prevalence of AD in America was estimated at 6.7 million in 2023, with projections suggesting a doubling to 13.8 million by 2060^2^. Despite extensive research in the last century, there remains no cure for AD, and current treatments are symptomatic rather than disease-modifying. With the increasing prevalence of AD driven by an aging population, there is an urgent need for continued research into its pathogenesis and the development of more effective therapeutic interventions.

Given these challenges, leveraging multi-omic data through advanced graph AI models presents a promising frontier in AD research. By integrating multi-omic data with graph-based techniques, researchers can more effectively identify critical disease biomarkers and uncover the core signaling pathways involved in AD. This approach offers the potential to not only enhance our understanding of AD pathogenesis but also pave the way for the development of targeted and disease-modifying treatments. In this study, we explore the application of a graph AI model on multi-omic datasets to identify key biomarkers and signaling interactions in AD, demonstrating its superiority in classification accuracy and its capability to highlight significant molecular mechanisms at various omic levels.

Recently, Graph Neural Networks (GNN) have gained prominence due to their capability to model relationships within graph-structured data^3–6^. And numerous studies have applied the GNN with the integration of the multi-omics data. MOGONET^7^ (Multi-Omics Graph cOnvolutional NETworks) initially creates similarity graphs among samples by leveraging each omics data, then employs a Graph Convolutional Network (GCN^3^) to learn a label distribution from each omics data independently. Subsequently, a cross-omics discovery tensor is implemented to refine the prediction by learning the dependency among multi-omics data. MoGCN^8^ adopts a similar approach by constructing a patient similarity network using multi-omics data and then using GCN to predict the cancer subtype of patients. GCN-SC^9^ utilizes a GCN to combine single-cell multi-omics data derived from varying sequencing methodologies. MOGCL^10^ takes this further by exploiting the potency of graph contrastive learning to pretrain the GCN on the multi-omics dataset, thereby achieving impressive results in downstream tasks with fine-tuning. Nevertheless, none of the aforementioned techniques contemplate incorporating structured signaling data like KEGG into the model. Moreover, general GNN models are limited by their expression power, i.e., the low-pass filtering or over-smoothing issues, which hampers their ability to incorporate many layers. The over-smoothing problem was firstly mentioned by extending the propagation layers in GCN^11^. Moreover, theoretical papers using Dirichlet energy showed diminished discriminative power by increasing the propagation layers^12^. And multiple attempts were made to compare the expressive power of the GCNs^13^, and it is shown that WL subtree kernel^14^ is insufficient for capturing the graph structure. Hence, to improve the expression powerful of GNN, the *K*-hop information of local substructure was considered in various recent research^13,15–19^. However, none of these studies was specifically designed to well integrate the biological regulatory network and provide the interpretation with important edges and nodes^20^. In this study, the unique and major contributions of this study are as follows: 1) developed a graph neural network (GNN) model for the mosGraphs, 2) analyzed multi-omic mosGraph datasets of AD, and 3) identified, visualized and evaluated a set of AD associated signaling biomarkers and network.

## Methodology and Materials

### Multi-omics datasets of Alzheimer’s Disease

To study Alzheimer’s Disease, multi-omics datasets were sourced from publicly accessible databases, specifically the ROSMAP datasets (refer to Table 1). Upon downloading these datasets, they were transformed into 2-dimensional data frames, structured with columns for sample IDs, sample names, etc., and rows for probes, gene symbols, gene IDs, etc. Integrating multi-omics data with clinical data necessitated identifying identical samples across the datasets. This process involved standardizing the rows (probes, gene symbols, gene IDs, etc.) into a uniform gene-level format, either by aggregating measurements for each gene or removing duplicates caused by gene synonyms. Genes were then aligned to a reference genome to ensure accurate final annotation in the multi-omics data. Standardization of gene counts across datasets was performed, and missing values were imputed with zeros or negative ones where necessary. Once all columns were aligned to standard sample IDs and all rows to standard gene IDs, and the number of samples and genes were unified, the data was ready for integration into Graph Neural Network (GNN) models. In these models, epigenomics, genomics, and transcriptomics data served as features for protein nodes.

**Table 1.**
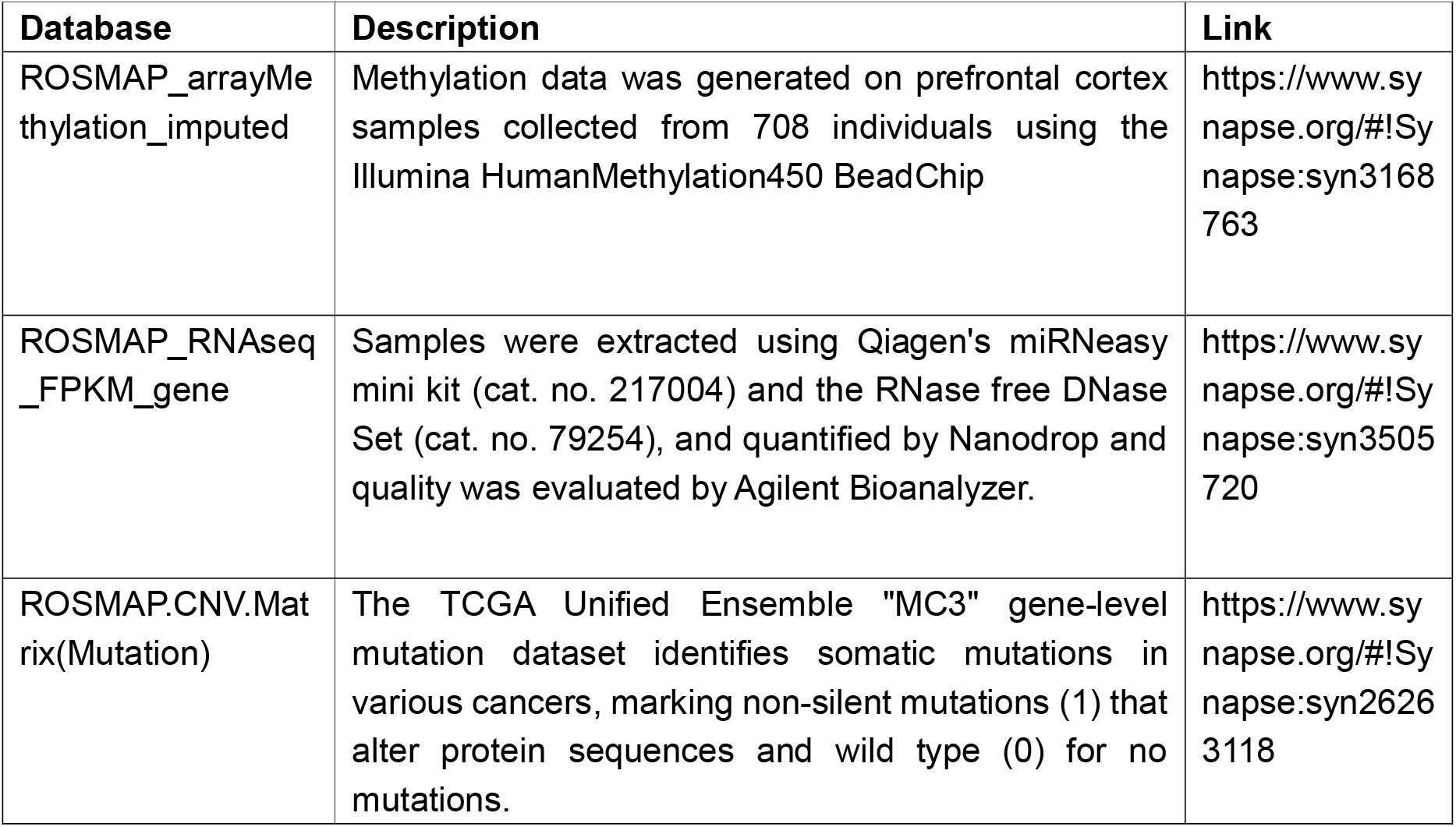
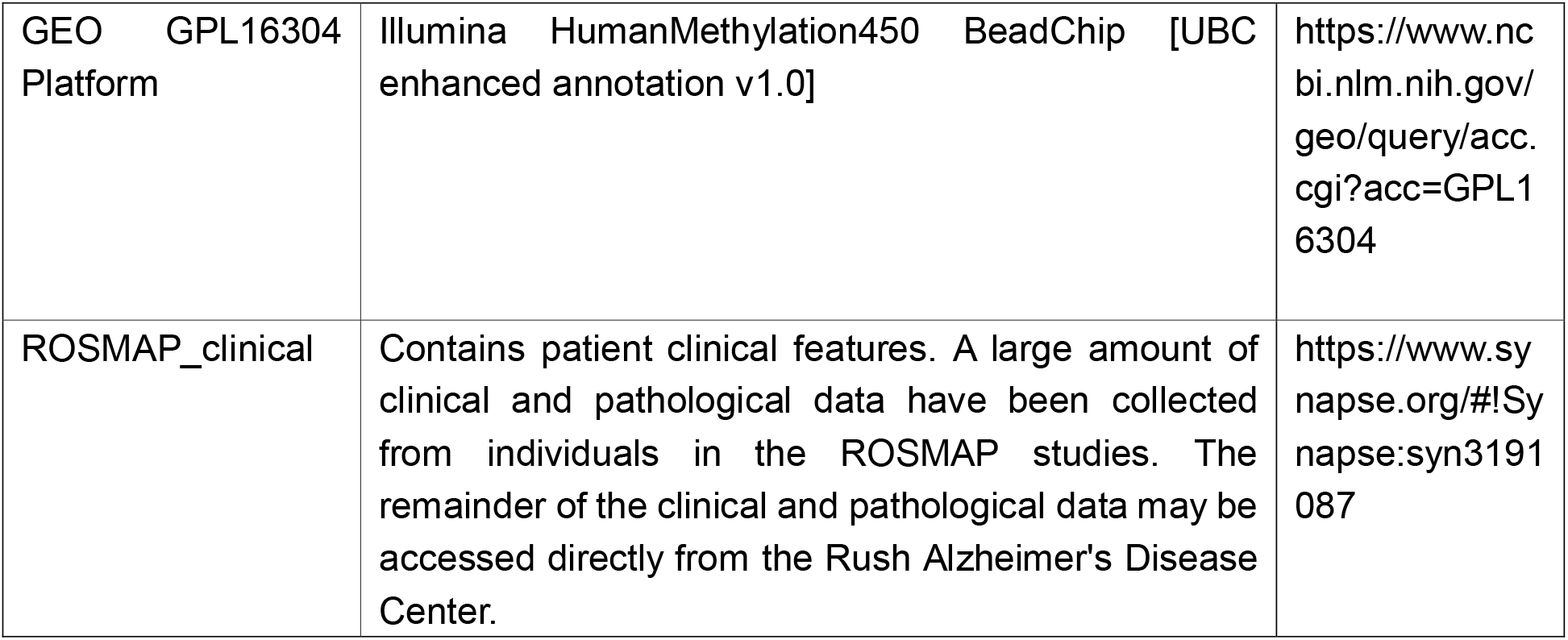
ROSMAP Database resources.

### KEGG Regulatory Network Construction

For constructing the knowledge graph, genes were selected by intersecting multi-omics datasets with gene regulatory networks from the KEGG database, which includes 2241 genes and 21041 edges. This intersection resulted in 2144 gene entities.

#### Architecture of the mosGraphFlow model

The proposed **mosGraphFlow** model enhances the analysis and prediction capabilities in multi-omics data, which aims to provide a comprehensive and interpretable analysis of AD dataset. The integrated approach offers a robust solution for multi-omics data analysis with generation of 𝒢=(*V,E)*, where |v| = n. In details, there are 3 types of nodes in the graph. *n*^(*meth*)^, *n*^(*gene*)^ and *n*^(*prot*)^ have the same number of nodes and *n*= *n*^(*meth*)^ + *n*^(*gene*)^ + *n*^(*prot*)^. Furthermore, the whole graph 𝒢 can be decomposed into subgraphs 𝒢 ′ and 𝒢 _*PPI*_, where 𝒢 ′ =𝒢 \ 𝒢 _*PPI*_ ; 𝒢 ′is the internal signaling flow graph which only contains the signaling flows from promoters to proteins; 𝒢_*PPI*_ is the protein-protein interaction (PPI) graph (|*V*_*PPI*_ | = *n*^(*prot*)^). Correspondingly, the adjacency matrices *A, A*′ and *A*_*PPI*_ for whole graph 𝒢, internal graph 𝒢 ′ and PPI graph 𝒢 _*PPI*_ will be generated. And the proposed model can be denoted as*f*(·), the graph-based deep learning model. It will predict the patient outcome *Y* (*Y* ∈ ℝ^*M*×1^), being constructed with *f*(𝒳,*A,A*_in_,*S*_*PPI*_)=*Y* where 𝒳= {*X*^(1)^,*X*^(2)^, …, *X*^(*m*)^,…, *X*^(*M*)^ } (*X*^(*m*)^ ∈ ℝ^*n*×*d*^)denotes all *M* data points in the dataset and *X*^(*m*)^ is the *m*-th data points. And *A*(*A* ∈ ℝ^*n*×*n*^) is the adjacency matrix that demonstrates the node-node interactions, and the element in adjacency matrix *A* such as *a*_*ij*_ indicates an edge from *i* to *j*.*A*′(*A* ′ ∈ ℝ^*n*×*n*^)is the adjacency matrix which only includes the node interactions from promoters to proteins, corresponding to the graph 𝒢 ′. Regarding the set of subgraphs in the PPI, 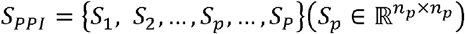, these subgraphs will partition the whole PPI graph adjacent matrix *A*_*PPI*_ into multiple subgraphs with the annotation of each individual signaling pathway, where the vertices in these partitioned subgraphs can be denoted as *V*_1_, *V*_2_,…, *V*_*p*_, …, *V*_*P*_, where 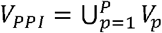. In each subgraph *S*_*P*_, there are nodes interactions between its internal *n*_*p*_ nodes and each subgraph has its own corresponding subgraph node feature matrix 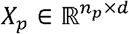.

#### Internal Modular Message Propagation

In the graph message passing stages of our architecture (see **Figure 1** step 2), we introduced the message passing between the internal links via matrix *A*′ with following formula:

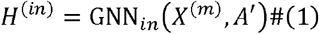

where *GNN*_*in*_ is the selected message propagation network and 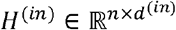 is the embedded node features after internal message propagation.

**Figure 1.**
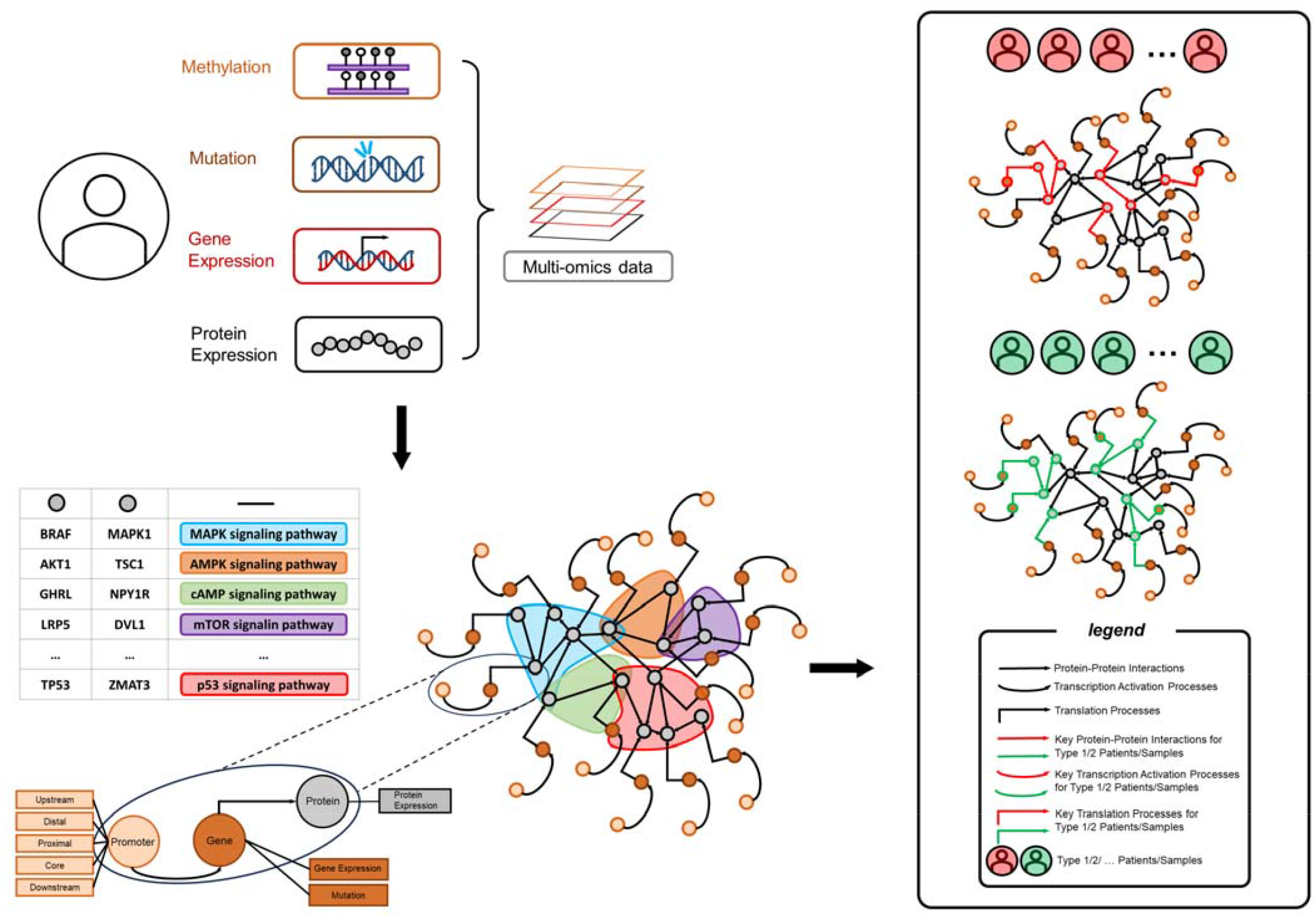
Architecture of ***mosGraphFlow***

#### Multi-hop Message Propagation in Signaling Pathway Subgraphs

Following the internal message passing stages, the local structure for each signaling pathway subgraph can be integrated via following formula:

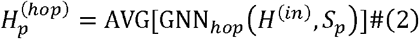

where *GNN*_*hop*_ is *K* -hop attention-based graph neural network borrowed from M3NetFlow^21^ framework and 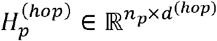.The aggregated node features 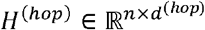 will be generated by AVG function for averaging the node included over multiple sub-signaling pathways in the set *s*_*PPI*_.

#### Global Bi-directional Message Propagation

Following the message propagation in the multiple internal subgraphs, the global weighted bi-directional message propagation^22^ will be performed via formula

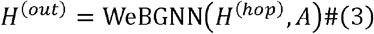

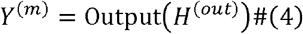

where WeBGNN (Weighted Bi-directional Graph Neural Network) is the graph signaling flow framework and 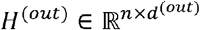. And linear transformation function 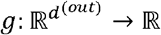 will be applied to outputting stage to predict the patient outcome with *Y*^(*m*)^.

## Results

### Experimental Settings

We utilized 437 samples from the ROSMAP dataset, categorized by disease status (275 AD, 162 non-AD) and gender (276 females, 161 males). Among the AD samples, there were 177 females and 98 males. To address the significant data imbalance, we performed downsampling for both classification tasks. For the AD vs. non-AD classification, we downsampled the AD samples to match the 162 non-AD samples, resulting in a dataset with 162 AD and 162 non-AD samples. For the gender classification within the AD samples, we downsampled the female AD samples to match the 98 male AD samples, resulting in a balanced dataset of 98 female and 98 male AD samples. We used 5-fold cross-validation to evaluate the performance of our models on both AD/non-AD and gender classification task.

### Model hyperparameters

The model was implemented using PyTorch and PyTorch Geometric, with the Adam optimizer employed for training. For the AD classification task, the initial learning rate was set to 0.002, and the training epochs were empirically set to 80. For the gender classification task within AD samples, the initial learning rate was set to 0.001, and the training epochs were set to 50. The hidden dimension was set to 10, and the leaky ReLU parameter was configured to 0.1. The output dimension was initially 30, which was subsequently reduced to 1 dimension through max pooling over the receptive field in the final pooling layer. A 5-fold cross-validation approach was utilized. The mean square error (MSE) and the correlation between the predicted gene effect scores and the experimental gene effect scores were used as the loss functions.

### Model performances and comparisons

Tables 2 and 4 present the accuracy and negative log likelihood (NLL) loss values for both the training and testing datasets, with **Table 2** displaying the values for AD/non-AD and **Table 4 for** female/male. The results indicate that the model achieved comparable performance on both datasets. Additionally, the proposed model was compared with other widely used models, namely GCN^23^, GAT^5^, GIN^6^, and UniMP^24^ (see **Table 3** and **Table 5**). The proposed model significantly outperformed the GAT, GCN, GIN, and UniMP models.

**Table 2.** NLL and accuracy values of the proposed model on the 5-fold cross-validation datasets (AD/non-AD)

| Number of folds | NLL on Training data | NLL on Testing data | Accuracy of Training data | Accuracy of Testing data |
| --- | --- | --- | --- | --- |
| 1<sup>st</sup> fold | 0.5443 | 0.5874 | 0.7192 | 0.6718 |
| 2<sup>nd</sup> fold | 0.5125 | 0.6913 | 0.7692 | 0.6563 |
| 3<sup>rd</sup> fold | 0.6309 | 0.6442 | 0.6423 | 0.6406 |
| 4<sup>th</sup> fold | 0.5696 | 0.6407 | 0.6961 | 0.6718 |
| 5<sup>th</sup> fold | 0.7248 | 0.6009 | 0.6931 | 0.5588 |

**Table 3.** Model comparison with other GNN network (AD/non-AD)

| Models | The Average NLL on Training data | The Average NLL on Testing data | The average accuracy of Training data | The average accuracy of Testing data |
| --- | --- | --- | --- | --- |
| Proposed model | 0.5964±0.0604 | 0.6329±0.0365 | 0.7039±0.0674 | 0.6398±0.0641 |
| GAT | 0.6649±0.0075 | 0.6935±0.0134 | 0.5942±0.0232 | 0.5710±0.0036 |
| GCN | 0.6753±0.0059 | 0.7824±0.1681 | 0.5734±0.0160 | 0.5553±0.0544 |
| GIN | 0.6756±0.0083 | 0.6832±0.0051 | 0.5469±0.0245 | 0.5377±0.0301 |
| UniMP | 0.6632±0.0029 | 0.8149±0.1583 | 0.6027±0.0271 | 0.5713±0.0219 |

**Table 4.** NLL and accuracy values of the proposed model on the 5-fold cross-validation datasets (female/male)

| Number of folds | NLL on Training data | NLL on Testing data | Accuracy of Training data | Accuracy of Testing data |
| --- | --- | --- | --- | --- |
| 1<sup>st</sup> fold | 0.6731 | 0.6734 | 0.7134 | 0.6667 |
| 2<sup>nd</sup> fold | 0.6884 | 0.6923 | 0.5987 | 0.5897 |
| 3<sup>rd</sup> fold | 0.6879 | 0.6892 | 0.6433 | 0.6410 |
| 4<sup>th</sup> fold | 0.6830 | 0.6869 | 0.6306 | 0.6154 |
| 5<sup>th</sup> fold | 0.6852 | 0.6937 | 0.6090 | 0.5500 |

**Table 5.** Model comparison with other GNN network (female/male)

| Models | The Average<br>NLL on Training<br>data | The Average<br>NLL on Testing<br>data | The average<br>accuracy of<br>Training data | The average<br>accuracy of<br>Testing data |
| --- | --- | --- | --- | --- |
| Proposed<br>model | 0.6835±0.0062 | 0.6871±0.0081 | 0.6390±0.0451 | 0.6126±0.0452 |
| GAT | 0.6790±0.0109 | 0.7369±0.0804 | 0.5880±0.0437 | 0.5562±0.0422 |
| GCN | 0.6753±0.0300 | 0.7553±0.0735 | 0.5549±0.0810 | 0.4586±0.1014 |
| GIN | 0.6963±0.0074 | 0.7296±0.0464 | 0.5051±0.0337 | 0.3924±0.0601 |
| UniMP | 0.6526±0.0220 | 0.6922±0.0488 | 0.6084±0.0254 | 0.5458±0.1070 |

### Signaling pathway inference

To interpret the underlying mechanisms of AD, the best-performing model was selected after training and validation process, then it was analyzed to extract attention scores from various graphs, which were used to infer signaling pathways related to the disease and key nodes (genes, promoters, and proteins). For each patient, the attention matrices of 1-hop neighbor nodes were calculated in every fold of the cross-validation process. Depending on the specific analysis, patients were stratified into different categories based on either their AD status (AD or non-AD) or gender (female or male), and for each category, the average attention matrices were computed. To quantitatively assess the significance of each node within these networks, the weighted degree of each node for every patient was calculated based on these attention scores, as detailed in the following formula:

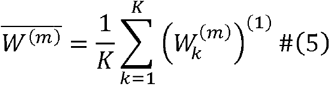

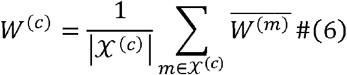

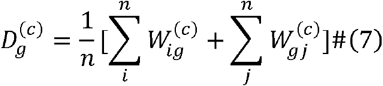

where 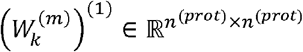 represents the attention-based matrix extracted from the first hop attention for patient *m* in the *k*-th fold; 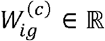 denotes the element of *K* -fold averaged attention matrix for patient *m* in the *i*-th row and *j*-th column for patient type 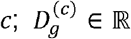 represents the node degree, which quantifies the importance of the node *g* within the network from the type of patient *c*.

Afterwards, the unimportant signaling flows in the attention-based matrix for certain type of patient will be filtered out by

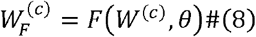

where *F* (·)is the filtering mapping function by providing selection of each element in the matrix with

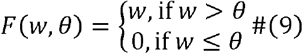

where *w* ∈ ℝ is the element in the input matrix and 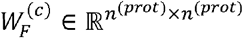 is the filtered matrix. Hence, the filtered node set for patient type 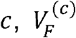,will be generated by removing independent nodes and nodes in those small connected components with number of nodes lower than *ϕ*, resulting in 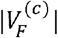 nodes.

### Sample-specific Network Visualizatio

The distinctions between AD/non-AD or female/male AD patients were made, and the relevant features for each group were identified. Subsequently, p-values for the gene features, such as methylations in promoter nodes, mutations and genes expression in gene nodes and proteins expression in protein nodes were calculated. The p-value calculation for these features was conducted by using the chi-squared test to check the differences between AD/non-AD samples or female/male of AD patients. This statistical method determined whether there were significant differences in the gene features between the samples of AD/non-AD or female/male from AD. By constructing contingency tables and performing the chi-squared test for each gene feature, p-values indicating the statistical significance of the observed differences were obtained. Ultimately, the top *T* gene features associated with AD or gender were selected based on these p-values.

After finalized important gene features ranked by p-values in top *T*, the network was pruned by iteratively removing the nodes which are only connected to one another unimportant node in a linear branch with node recursive algorithm (check details of this algorithm in **Appendix A.1** and **Figure S1**). This ensures that each remaining nodes is either linked to an important node or is part of a more complex interaction network, enhancing the purity and reliability of the gene interaction data.

Subsequently, nodes degree were calculated to identify hub node (node degree larger than 2). The set of middle nodes for certain path *t* which connects two hub nodes *u* and *ν* can be denoted as 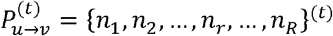, where *λ* + 1 is the length of path. Hence, the average edge weight on the path 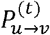 can be generated by

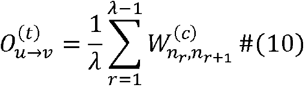

where 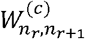 is the edge weight from node *n*_*r*_ to node *n*_*r* +1_. For all of the paths detected between the hub node *u* and hub node *ν*, the nodes on the top *β* paths will be kept. Additionally, p-value middle nodes, which are crucial due to their statistical significance, will be retained along with middle nodes that are adjacent to these p-value nodes. (check **Appendix Section A.2** for details).

### Inferred core signaling networks of selected patient type

By setting an edge threshold *θ* as 0.12, and a small component threshold *ϕ* as 20, we identified 183 and 175 potential important protein nodes for AD and NOAD, respectively. Then, by calculating the p-value < 0.2, the top 70 gene features associated with Alzheimer’s Disease were selected. By pruning linear branches and keeping the nodes via top 2 (*β* = 2) paths between hub nodes, we filtered out non-essential nodes, reducing the number of potential important protein nodes to 152 for AD and 136 for non-AD. The corresponding gene weights 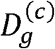 for these top 70 (*T*= 70) gene features for AD/non-AD were calculated. These top 70 gene features and gene weights are shown in **Figure 4** and detailed in **Table 6**. In **Figure 2** and **Figure 3**, these node from top gene features (promoters, transcriptions and mutations) are represented by non-blue nodes derived from the blue nodes, with different colors indicating various types of gene features. Different sizes of the nodes represent the varying importance of the gene features, with larger nodes indicating greater significance based on their p-values.

**Table 6.**
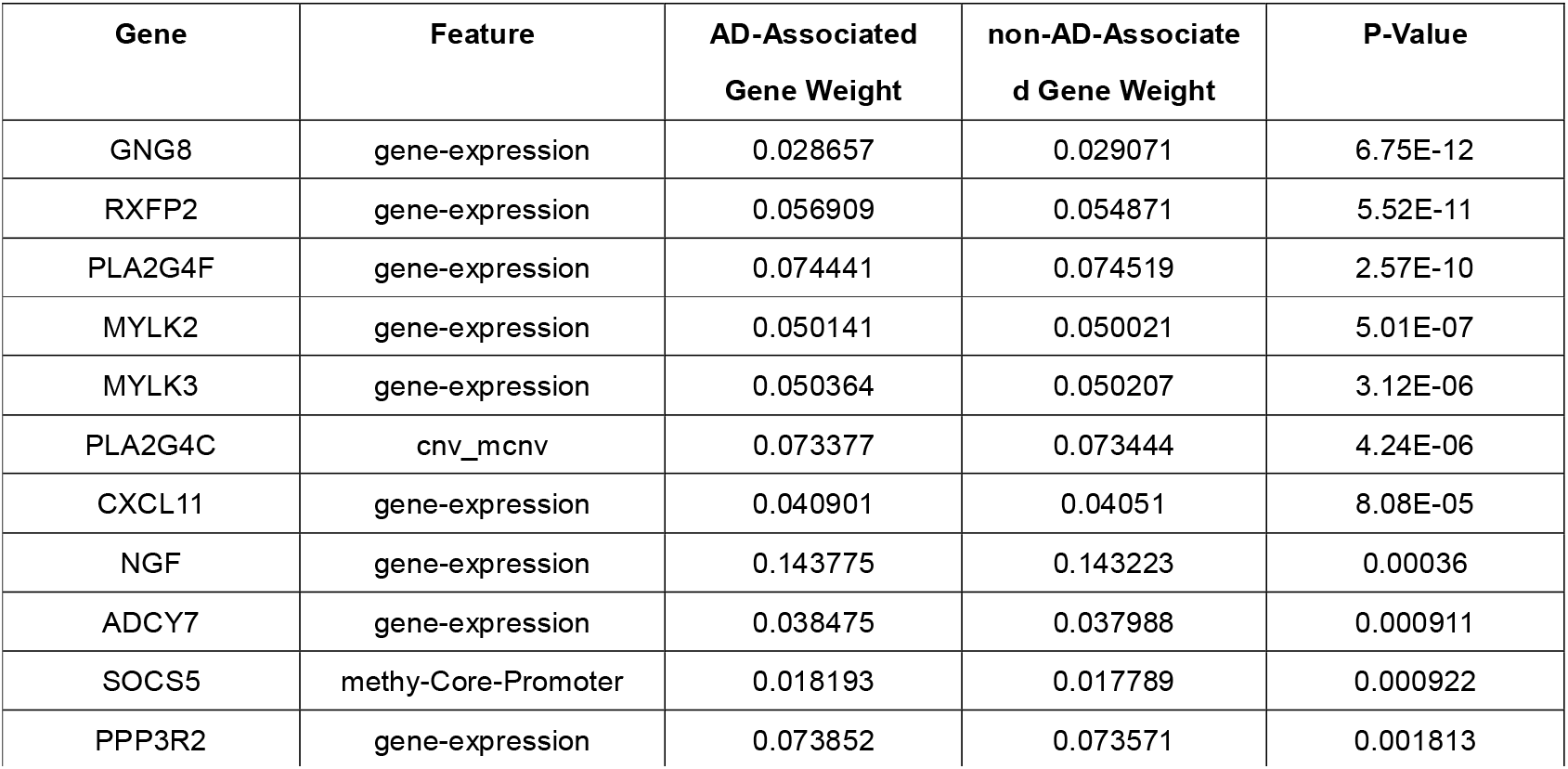
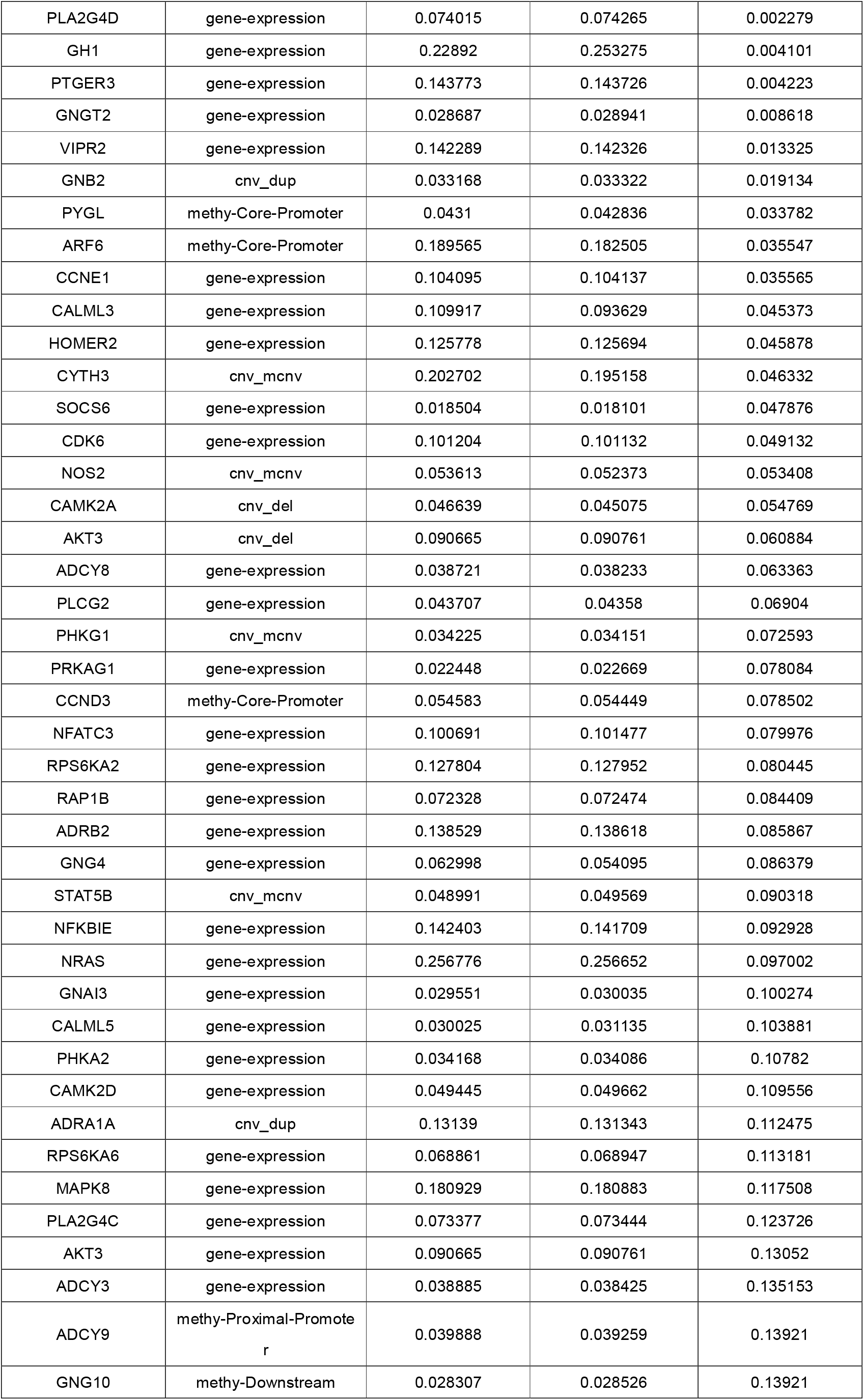
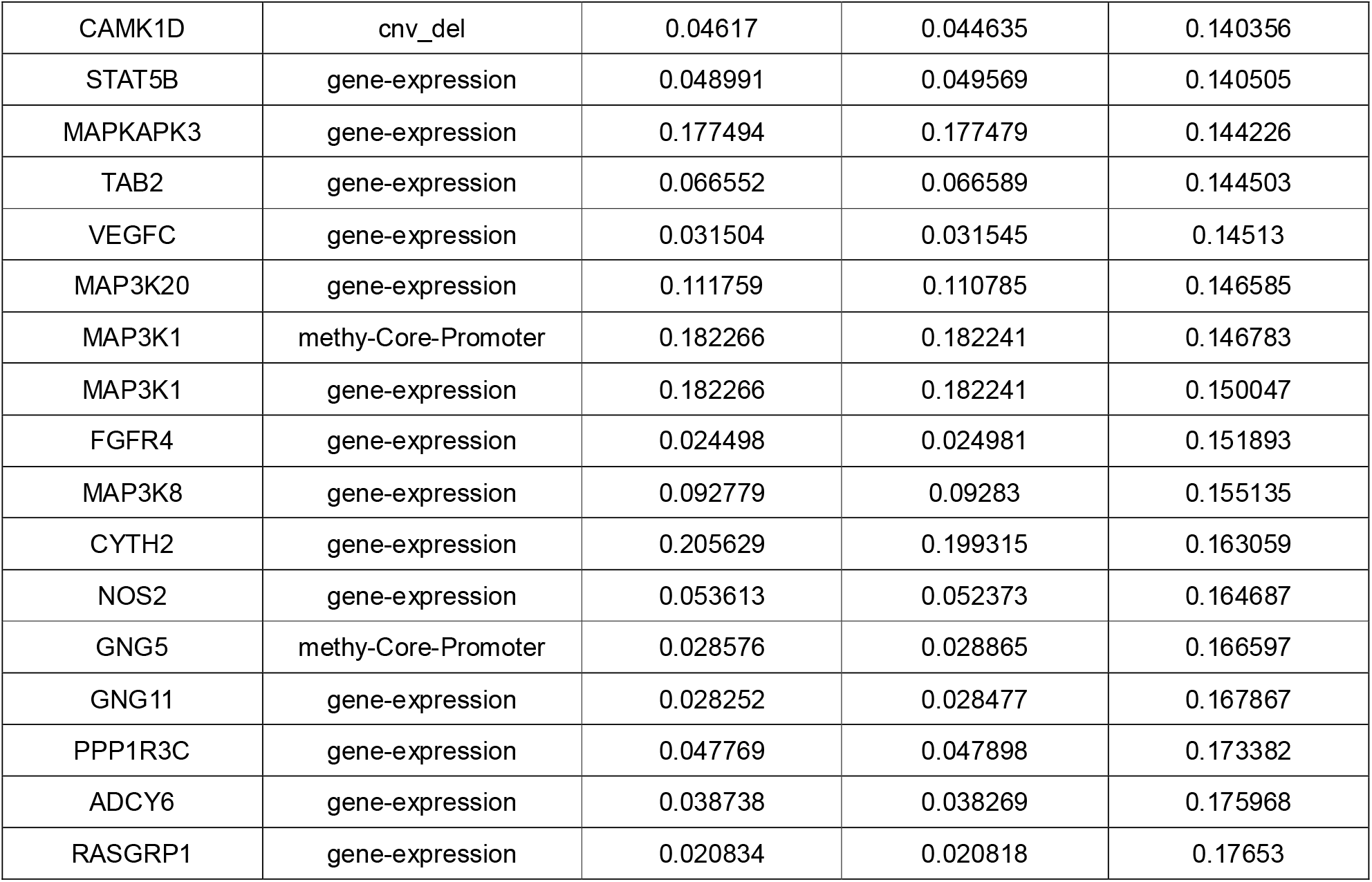
Top 70 gene features associated with Alzheimer’s Disease

**Figure 2.**
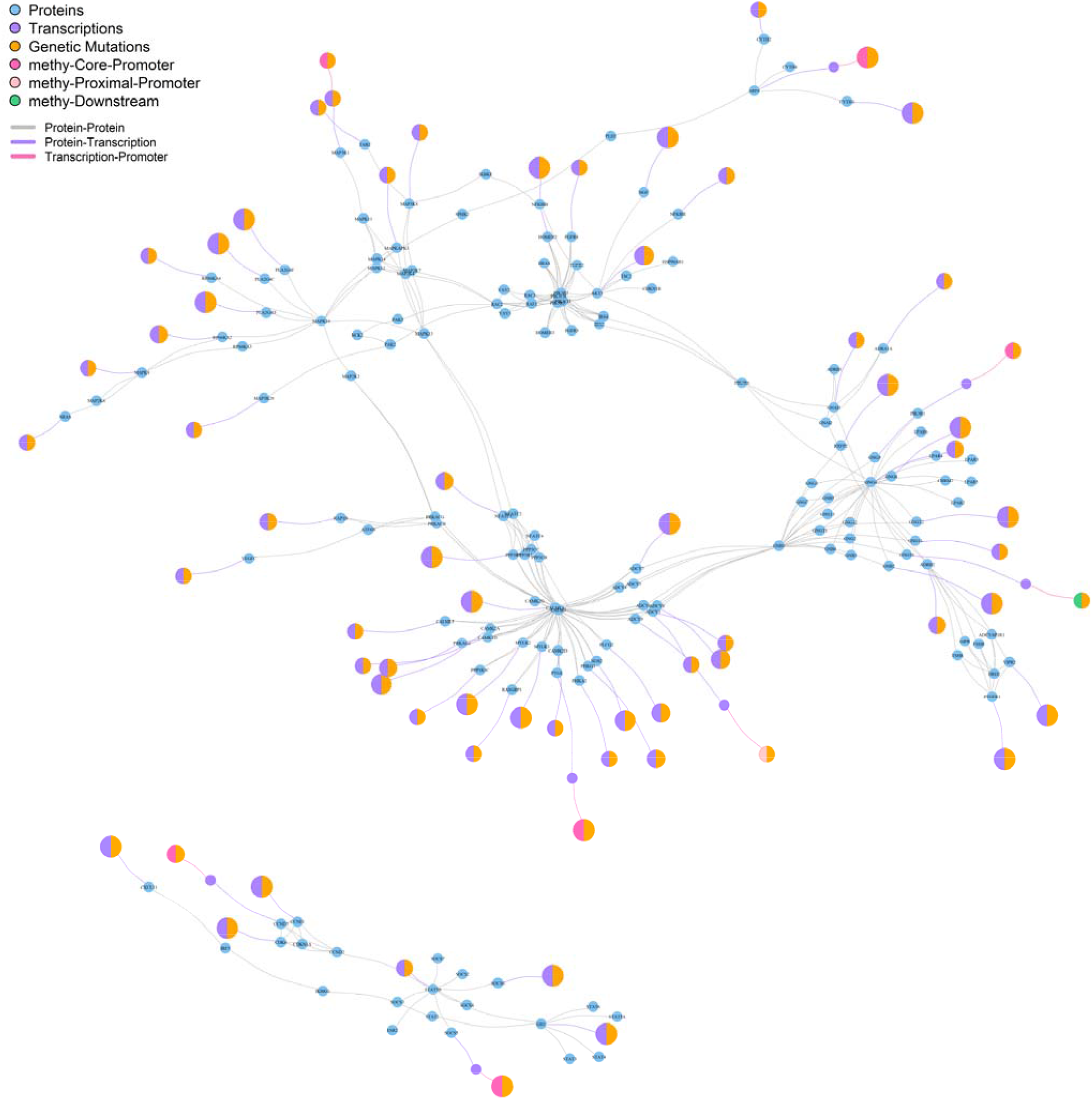
Top 70 important nodes signaling network interaction in AD samples

**Figure 3.**
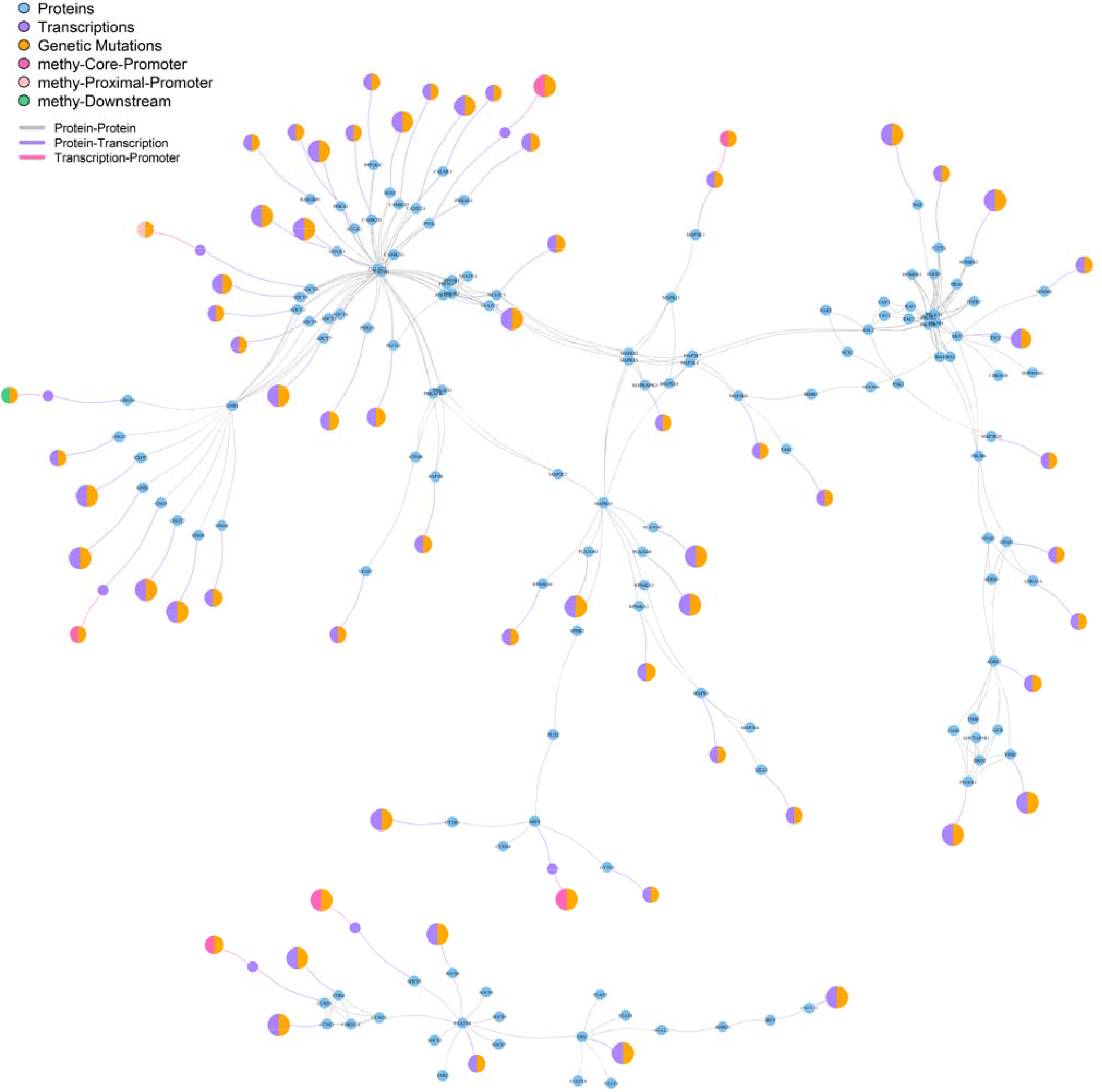
Top 70 important nodes signaling network interaction in non-AD samples

**Figure 4.**
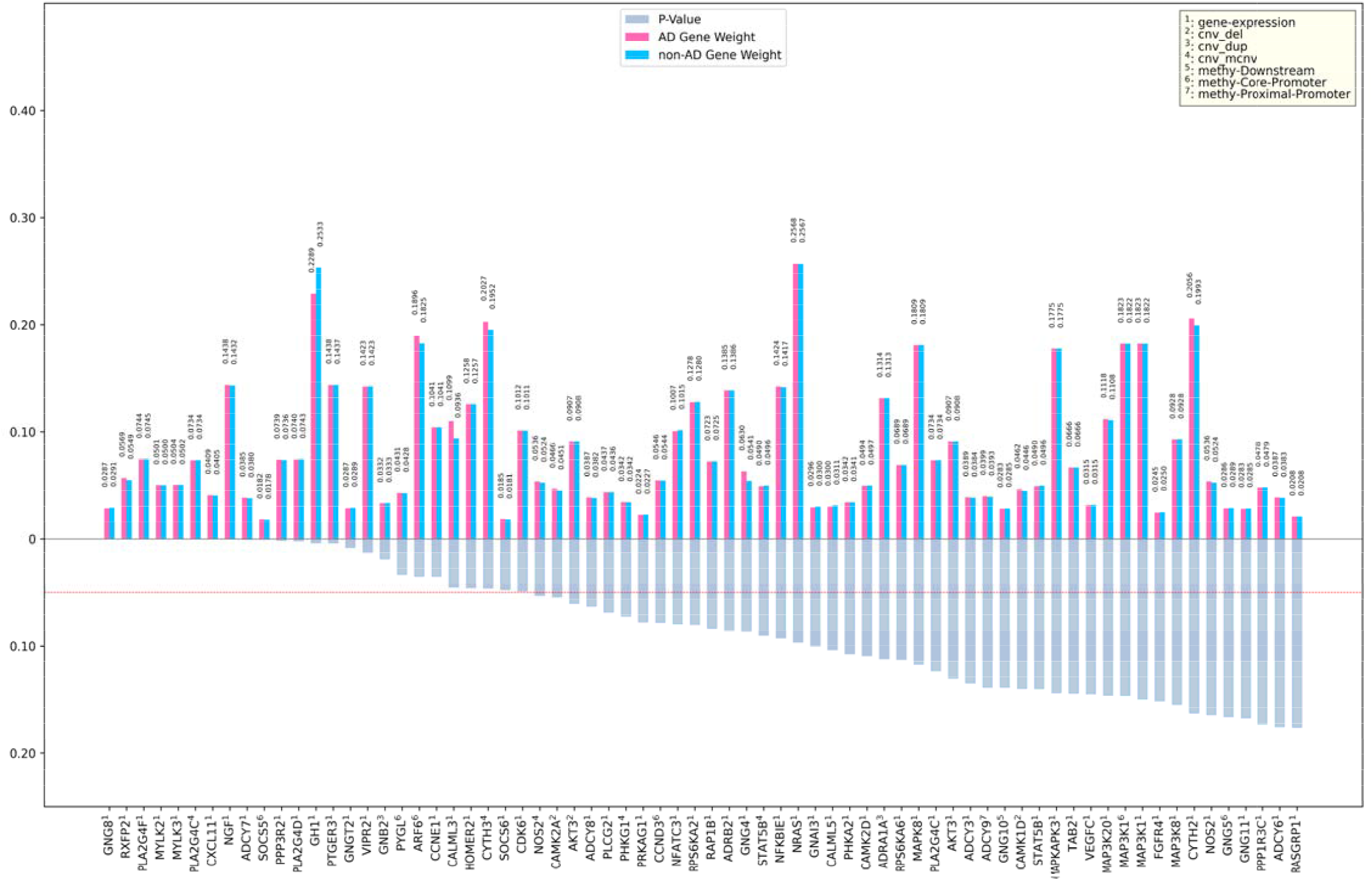
Bar chart displaying the weight of important genes in AD and non-AD samples, ranking by their p-values. (The red dashed line indicates a p-value threshold of 0.05)

Similarly, **Figure 5** and **Figure 6** shows the inferred core signaling networks with top 70 gene features for female and male subjects, and these the top 70 gene features and gene weights are shown in **Figure 7**. In this analysis, through node optimization process, similar to the above, we identified 214 potential important protein nodes for females and 214 for males. Notably, we observed a significant overlap between the protein nodes selected from the AD signaling networks and those from the female signaling networks. Specifically, there are 81 overlapping protein nodes between the 152 protein nodes identified in the AD signaling networks and the 214 protein nodes identified in the female signaling networks (see **Appendix B Table S1**). Furthermore, there is an overlap of 15 gene features between the top 70 AD/non-AD gene features and the top 70 female/male gene features (see **Appendix B Table S2**). This overlap further supports the feasibility of our proposed model in identifying key target genes for AD.

**Figure 5.**
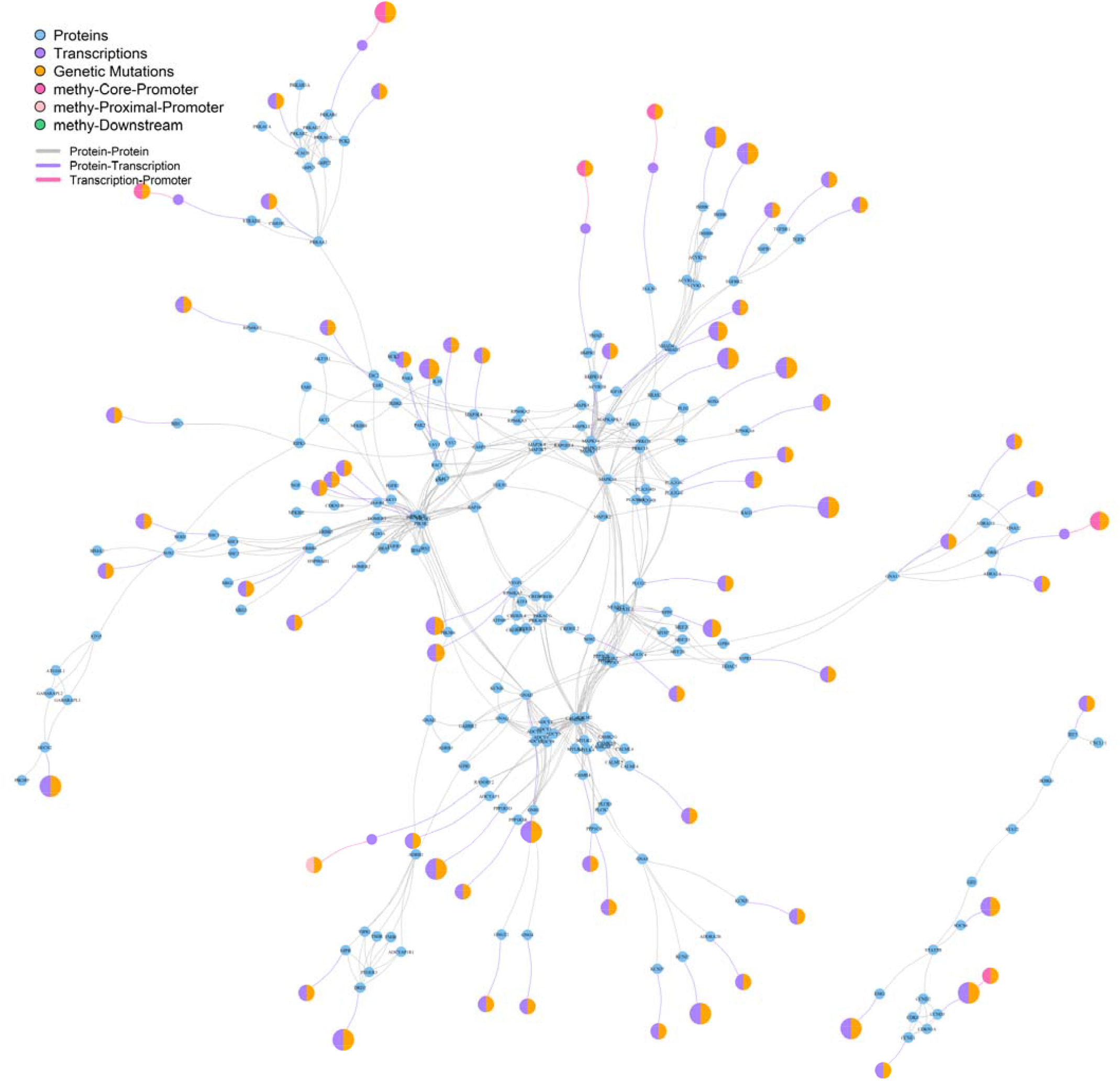
Top 70 important nodes signaling network interaction in females

**Figure 6.**
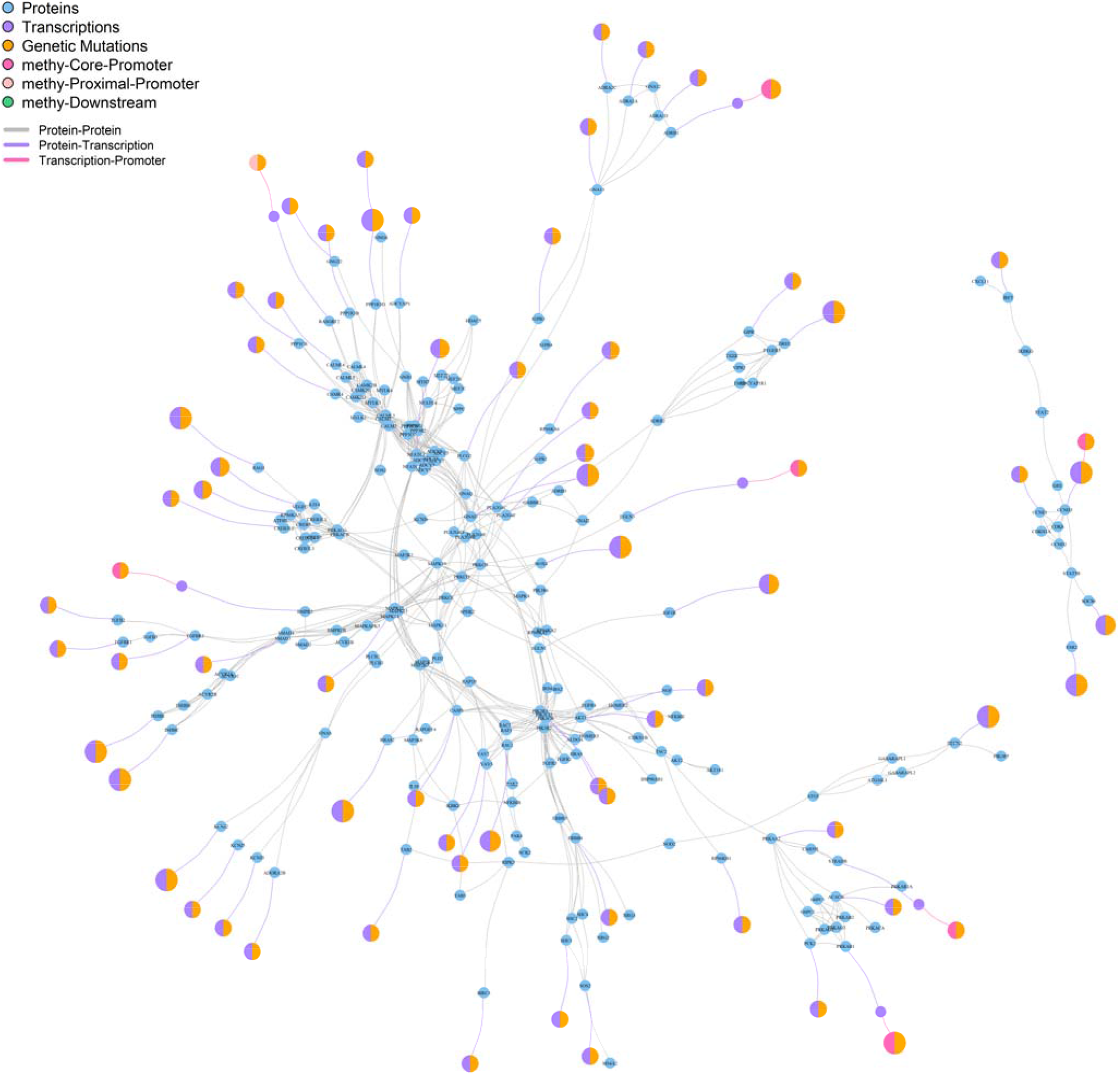
Top 70 important nodes signaling network interaction in males

**Figure 7.**
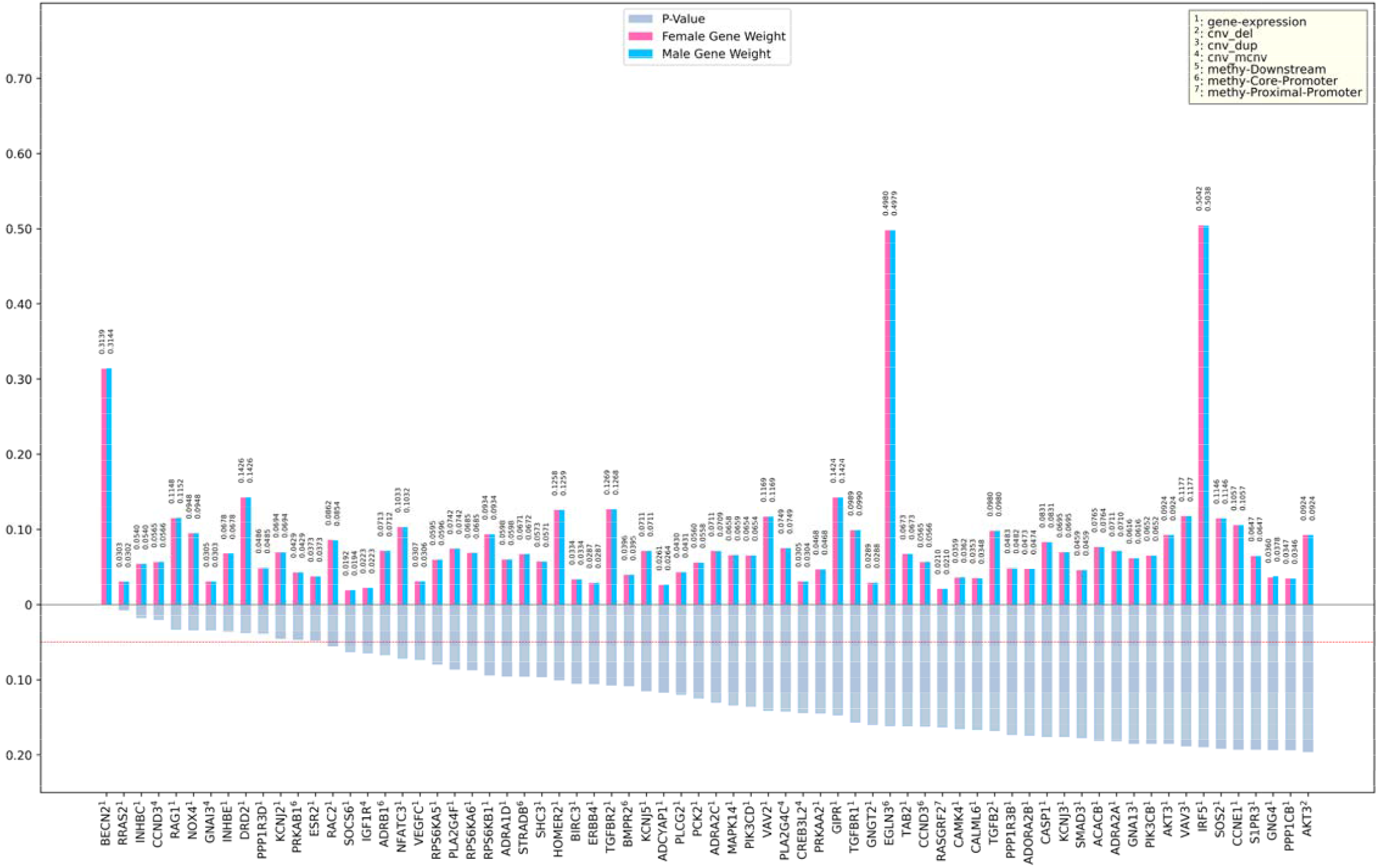
Bar chart displaying the weight of important genes in female and male, ranking by their p-values. (The red dashed line indicates a p-value threshold of 0.05)

### Model validation: pathway enrichment analysis

Pathway enrichment analysis was conducted on the top 70 genes associated with AD using ShinyGO 0.80 and the KEGG pathway database. This analysis revealed the top 20 signaling pathways involving these genes (see **Figure 8**), enhancing our biological understanding of their roles in AD pathogenesis.

**Figure 8.**
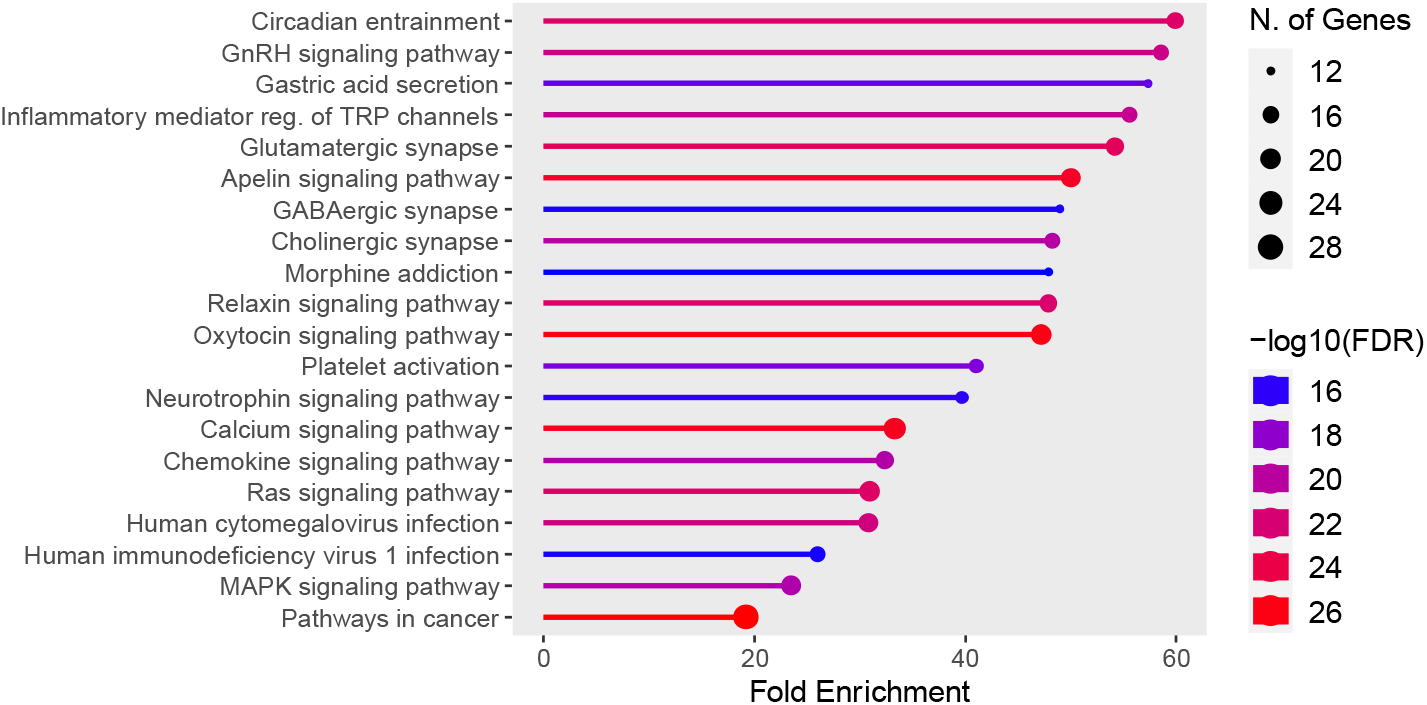
Lollipop plot showing the negative base-10 logarithm of the False Discovery Rate (FDR) and number of genes of the top 20 signaling pathways based on the top 70 gene features found to be associated with AD. Generated by ShinyGo 0.80 after performing pathway enrichment analysis with FDR cutoff at 0.05.

To gain a comprehensive view of the complex nature of these signaling pathways, we utilized a Sankey diagram to visualize the interconnectedness between the top 70 genes and their associated pathways (see **Figure 9**). The KEGG pathway database categorizes signaling pathways into seven broad categories: metabolism, genetic information processing, environmental information processing, cellular processes, organismal systems, human diseases, and drug development. However, for a more detailed focus on function or disease-related aspects, specific categories are highlighted (see **Figure 9** and **Appendix C Table S3**).

**Figure 9.**
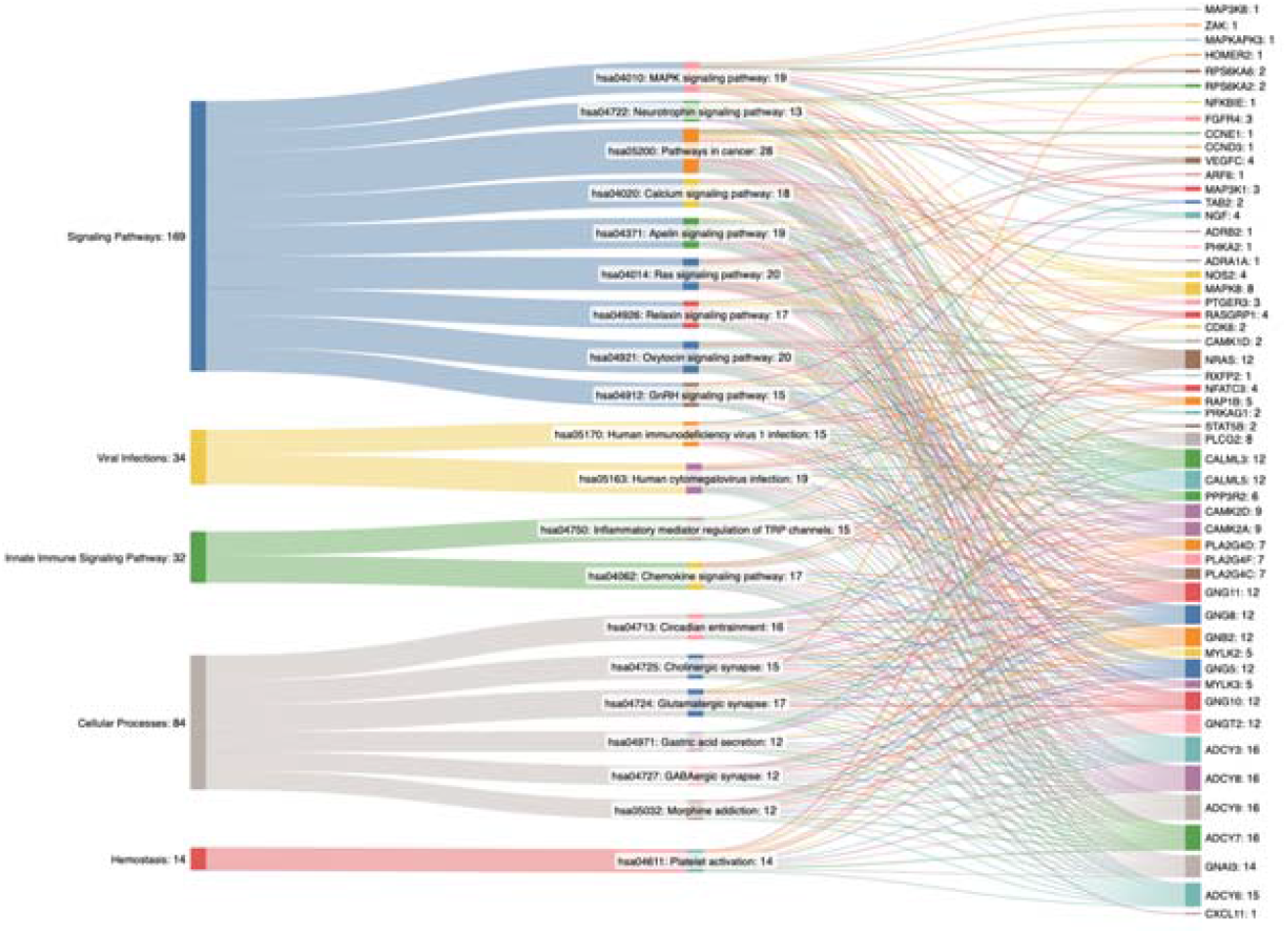
Sankey diagram illustrating the relationship between the identified signaling pathways and corresponding genes using the top 70 genes features found to be associated with AD

### Model validation: gene validation

The identification of the top 20 genes associated with AD in females and males, listed in **Table 8**, represents a significant step forward in understanding the genetic underpinnings of this condition. The genes were ranked using our novel graph AI model, which integrated multi-omic data to highlight key biomarkers and signaling interactions relevant to AD. Notably, the same genes were identified for both sexes, underscoring their critical role in AD pathogenesis. **Table 8** provides a comprehensive overview of these genes, detailing their functions and specific relations to AD. The function of each gene and its contribution to AD pathology are pivotal for elucidating the complex biological mechanisms driving the disease. The consistency of gene rankings between females and males emphasizes the universal importance of these biomarkers in AD. This uniformity suggests that the identified genes play fundamental roles in the disease’s progression, regardless of sex. This finding enhances the potential for developing targeted therapies that could benefit a broad patient population.

**Table 8.** Top 20 genes associated with Alzheimer’s Disease in females and males

| Gene | Function and Relation to AD |
| --- | --- |
| RRAS2 | RRAS2 is a key regulator of G-protein-coupled receptor signaling and neuronal plasticity. Reduced expression of RRAS2 correlates with cognitive decline in AD patients<sup>76</sup>. In AD mouse models, RRAS2 modulates neuronal hyperactivity, memory impairment, dendritic spine loss, and neuronal cell death<sup>77</sup>. Notably, Aβ-induced neuronal hyperactivity can be mitigated by targeting the ryanodine receptor 2 (RyR2) to reduce its open time<sup>78</sup>. Neuronal hyperactivity is an early defect observed in both familial and sporadic AD, accelerating the onset of neuronal dysfunction<sup>79</sup>. Given this, it is worthwhile to investigate the potential relationship between RRAS2 and RyR2. Understanding this interaction could lead to the development of novel therapeutics aimed at reducing neuronal hyperactivity and its associated neurodegenerative consequences, ultimately improving outcomes for AD patients. |
| RAG1 | RAG-1 is vital for recombining immunoglobulin and T-cell receptor genes, essential for developing mature B and T lymphocytes, underscoring its critical immune role<sup>80</sup>. RAG-1 is also expressed in the brain, especially in high neural density regions like |
|  | the hippocampus, though its precise brain function is unclear<sup>81</sup>. Research shows that hippocampal RAG-1 knockdown impairs spatial learning and memory in rats, suggesting a cognitive role<sup>81</sup>. Additionally, studies found that RAG-1, but not RAG-2, deficiency impairs social recognition memory, indicating a specific memory function<sup>80</sup>. Pathway analysis reveals that RAG-1 absence inhibits signaling pathways controlling neurodegenerative disorders, including AD, supporting the view of these diseases as having an autoimmune component<sup>80</sup>. |
| **PPP1R3D** | N/A |
| **RAC2** | In AD, synaptic dysfunction and loss are key contributors to cognitive deficits, with RAC2 implicated in these processes. Studies have shown that RAC2 is dysregulated in AD patients, with altered expression in the brain, cerebrospinal fluid, and blood<sup>82</sup>. This dysregulation is linked to the disruption of the actin cytoskeleton, crucial for maintaining dendritic spine stability and synaptic plasticity. RAC2 and other actin-binding proteins regulate the actin cytoskeleton's dynamics, essential for dendritic spine morphology<sup>82</sup>. Furthermore, RAC2 is involved in the Rho GTPase pathway, which is deregulated in several neurodegenerative disorders, including AD. The translocation of Rho GTPases to neurofibrillary tangles in dystrophic neurites correlates with the neuronal dystrophy observed in AD, and RAC2 activity is crucial here<sup>83</sup>. Additionally, RAC2 mediates dendritic degeneration induced by the cleavage of g-adducin, a cytoskeleton-associated protein, by asparagine endopeptidase. This cleavage disrupts the spectrin-actin assembly, leading to defects in neurite outgrowth and contributing to AD-like pathology and cognitive deficits in transgenic mouse models<sup>84</sup>. The protective effects of treatments like acetylcholinesterase inhibitors, which mitigate RAC2-related damage by regulating neurite outgrowth-related genes, further underscore RAC2's role in AD pathology. |
| **NFATc3** | NFATc3, a member of the NFAT family, regulates the transcription of genes involved in T-cell activation and angiogenesis. It is essential for the expression of IL2 and COX2 in T cells, crucial for T-cell proliferation and inflammatory responses<sup>85</sup>. NFATc3 also regulates COX2 expression in endothelial cells, necessary for COX2-dependent migration and angiogenesis in vivo<sup>85</sup>. Given the involvement of inflammation and vascular dysfunction in AD, NFATc3 could potentially play a significant role in this neurodegenerative disorder. Chronic inflammation is a hallmark of AD, and the up-regulation of inflammatory mediators like IL2 and COX2 could exacerbate neuronal damage and cognitive decline. Impaired angiogenesis and endothelial cell function are also implicated in AD, contributing to reduced cerebral blood flow and the breakdown of the blood-brain barrier. Dysregulation of NFATc3 could lead to increased inflammation and vascular abnormalities, both critical factors in AD development and progression. However, a direct role of NFATc3 in AD has not yet been established. |
| **PLA2G4F** | N/A |
| **RPS6KA6** | RPS6KA6 is a serine-threonine kinase in the MAPK pathway, regulating cell proliferation, survival, growth, and movement. In AD, RPS6KA6 is linked to neuroinflammation and synaptic damage, key pathological features. Dysregulation of RPS6KA6 affects the neurotrophin pathway, crucial for neuronal survival, neuroplasticity, and neurogenesis<sup>86</sup>. Neurotrophins maintain neuronal health, and their dysregulation can lead to neurodegeneration and cognitive decline in AD. RPS6KA6 also regulates long-term potentiation and axon guidance, vital for learning and memory, suggesting that its dysregulation impacts synaptic plasticity, contributing to AD-related cognitive deficits<sup>86</sup>. Autoantibodies against RPS6KA6 have been identified as potential biomarkers for predicting age-related neurodegenerative diseases, including AD and advanced-stage Parkinson's disease<sup>86</sup>. Inhibitors of ribosomal s6 kinase (RSK) signaling have shown promise in preventing seizures with elevated RSK activity and normalizing dendritic spine density in mouse models of neurodevelopmental disorders, indicating potential therapeutic strategies for AD<sup>87</sup>. |
| **PLCG2** | PLCG2 significantly impacts AD pathology through its role in inflammation and microglial function. Research shows PLCG2 expression is upregulated in several brain regions of late-onset AD patients and correlates with amyloid plaque density and microglial markers AIF1 and TMEM119<sup>88</sup>. This upregulation is also seen in the 5xFAD mouse model of AD, where PLCG2 expression increases with disease progression, especially in microglia<sup>88</sup>. The protective PLCG2 P522R variant has a slight hypermorphic effect on enzyme function, suggesting therapeutic benefits from activation rather than inhibition of PLCG2<sup>89</sup>. This variant also recruits CD8+ T cells to the brain, enhancing microglial antigen presentation and T cell recruitment, contributing to a protective microglial transcriptional state<sup>90</sup>. Additionally, the rs72824905-G allele in PLCG2 is linked to a reduced risk of AD, frontotemporal dementia, and dementia with Lewy bodies, indicating overlapping PLCG2-related immune signaling pathways in these diseases<sup>91</sup>. Furthermore, as a downstream component of TREM2 signaling, PLCG2 promotes survival, proliferation, phagocytosis, and cytokine secretion, all crucial for neurodegenerative processes<sup>89,90</sup>. |
| **MAPK14** | One of the primary roles of MAPK14 in AD is its involvement in the autophagic-lysosomal pathway, which is crucial for the degradation of misfolded proteins. In healthy neurons, low levels of MAPK14 ensure proper autophagic flux and degradation of BACE1, a protein involved in amyloid plaque formation. However, in AD neurons, increased levels of MAPK14 impair autophagic-lysosomal protein degradation, leading to higher BACE1 levels and subsequent plaque formation. Reducing MAPK14 levels has been shown to suppress these autophagic defects, thereby reducing BACE1 levels and plaque formation, highlighting its potential as a |
|  | therapeutic target<sup>92</sup>. Additionally, MAPK14 activation is linked to the hyperphosphorylation of tau proteins, another hallmark of AD. Extracellular Aβ accumulation triggers intracellular MAPK14 activation, which in turn leads to tau hyperphosphorylation and further Ab accumulation, creating a vicious cycle that exacerbates AD pathology<sup>93</sup>. |
| **PIK3CD & AKT3** | PIK3CD encodes the catalytic subunits for the PI3K heterodimer. In AD, the PI3K/Akt pathway is significantly implicated in forming NFTs and amyloid plaques, key disease features. PI3K activation produces PIP3, which recruits Akt to the cell membrane, where PDK1 and mTORC2 phosphorylate and activate it<sup>94,95</sup>. This cascade regulates downstream targets like GSK-3β, a key player in tau phosphorylation and subsequent NFT formation<sup>95,96</sup>. PI3K/Akt signaling also regulates Aβ production and deposition. Impaired PI3K/Akt signaling increases GSK-3β activity, enhancing Aβ production and aggregation, worsening AD pathology. Additionally, AD brains show decreased phosphorylation of PI3K/Akt pathway components, correlating with reduced insulin signaling and increased oxidative stress, which contribute to neuronal apoptosis and cognitive decline<sup>95,97</sup>. This pathway influences mitochondrial function and oxidative stress responses, essential for neuronal health. Dysregulation leads to increased oxidative stress and mitochondrial dysfunction, promoting neurodegeneration. |
| **VAV2** | VAV2, a guanine nucleotide exchange factor for Rho family GTPases, significantly impacts AD pathology through its interaction with the amyloid precursor protein (APP). APP is a crucial transmembrane protein in AD pathogenesis, and the tyrosine phosphorylation site Y682 on its intracellular tail is essential for its function. VAV2 directly binds to the Y682-phosphorylated APP tail via its SH2 domain, inhibiting APP degradation and increasing the levels of APP and its cleavage products, which are implicated in AD<sup>98</sup>. VAV2 is also involved in various biological processes, including endothelial cell function, vasodilation, blood pressure regulation, and neurogenesis, indicating its importance in early brain development<sup>99</sup>. VAV2 regulates neurite outgrowth and branching and interacts with TrkB receptors upon BDNF stimulation, activating Rac1, which is crucial for synaptic development and plasticity<sup>100</sup>. Nonetheless, the interaction between VAV2 and APP, particularly the stabilization of APP and its cleavage products, suggests a potential mechanism by which VAV2 contributes to Aβ accumulation. |
| **PLA2G4C** | N/A |
| **GNGT2** | Research has shown that GNGT2, along with ABI3, is part of a tightly co-expressed gene network in both AD patients and mouse models, indicating a potential collaborative role in AD pathogenesis<sup>101</sup>. The deletion of GNGT2, particularly in conjunction with ABI3, has been observed to influence AD pathology in distinct ways. |
|  | For instance, in the TgCRND8 mouse model, the loss of ABI3 and GNGT2 resulted in a significant reduction in amyloid plaque numbers and size, suggesting that these genes may modulate microglial behavior to enhance plaque clearance. This reduction in plaque pathology was accompanied by a decreased number of microglia clustering around the plaques, which implies that GNGT2 might be involved in the microglial response to amyloid deposition<sup>102</sup>. Additionally, transcriptomic analyses have revealed that the loss of GNGT2 leads to the upregulation of several AD-associated neurodegenerative markers, such as Trem2, Plcg2, and Tyrobp, even in the absence of typical AD neuropathology, further supporting its role in the disease's molecular mechanisms<sup>101</sup>. Systems biology approaches have also highlighted alterations in energy metabolism and the depletion of neuroprotective metabolites in AD, which could be linked to the dysregulation of genes like GNGT2<sup>103</sup>. |
| **TAB2** | TAB2, expressed in the brain and vascular endothelium, is suggested to regulate NFkB-mediated gene expression, which is crucial for inflammatory responses<sup>104</sup>. Recent studies highlight the potential of long non-coding RNAs (lncRNAs) in diagnosing and understanding AD. Exosomes, extracellular nanovesicles involved in immune response and neuronal function, carry lncRNAs differentially expressed in AD patients. Advanced bioinformatics and machine learning have identified specific lncRNAs, such as ENST00000608936 and ENST00000433747, as promising diagnostic markers for AD, with high sensitivity, specificity, and accuracy<sup>105</sup>. The interplay between TAB2's role in inflammation and the emerging significance of lncRNAs in AD suggests that TAB2 is a promising target for further investigation to gain deeper insights into the disease's pathology. |
| **CAMK4** | CAMK4 plays a significant role in the pathology of AD through its involvement in the hyperphosphorylation of tau protein and formation of NFTs, which are associated with neuronal death and cognitive dysfunction in AD patients<sup>106,107</sup>. The activation of CaMK4 is closely linked to calcium homeostasis within neurons. Elevated intracellular calcium levels, often resulting from Aβ deposition, activate the CaM-CaMK4 signaling pathway, leading to the phosphorylation of tau protein<sup>106</sup>. This process is exacerbated by the presence of familial Alzheimer's disease (FAD) mutant presenilins, which cause constitutive activation of CaMK4 and the transcription factor CREB, further promoting tau hyperphosphorylation and neuronal death<sup>108</sup>. Additionally, studies have shown that inhibiting the CaMKK-CaMK4 pathway can prevent the hyperphosphorylation of synapsin and CaMK4, suggesting that this pathway is crucial in the synaptic dysfunction observed in AD<sup>109</sup>. The neuroprotective effects of certain compounds, such as genistein and chrysophanol, have been attributed to their ability to modulate the CaM-CaMK4 pathway, thereby reducing tau hyperphosphorylation and offering potential therapeutic avenues for AD<sup>106,107</sup>. |
| **PPP1R3B** | PPP1R3B plays a role in glycogen metabolism and is associated with high-density |
|  | plasma lipoproteins, which previously has been documented to affect the clearance of β-amyloid in AD mouse brains<sup>110</sup>. Given the connection between lipid metabolism and AD, enhancing PPP1R3B activity could potentially improve lipid clearance pathways and reduce amyloid burden. This hypothesis is supported by findings that genetic variants influencing tau deposition, such as those in PPP2R2B, also affect metabolic pathways, suggesting a broader interplay between metabolism and neurodegeneration<sup>111</sup>. Therefore, targeting PPP1R3B to enhance glycogen metabolism and lipid clearance could be a novel therapeutic approach to mitigate amyloid accumulation in AD. |
| **PIK3CB** | PIK3CB is significantly downregulated in AD patients compared to controls, suggesting its potential involvement in the disease's progression<sup>112</sup>. The PI3K/AKT pathway, in which PIK3CB is a crucial component, is essential for cell survival, autophagy, neurogenesis, neuronal proliferation, differentiation, and synaptic plasticity, all of which are critical for maintaining cognitive function and neuronal health<sup>113</sup>. In Jnk3 null mice, an increase in PIK3CB transcription and protein levels was observed, leading to enhanced PI3K activity and AKT phosphorylation, which are associated with neuroresistance to cell death<sup>114</sup>. This suggests that PIK3CB may have a protective role against neurodegenerative processes. Additionally, the dysregulation of calcium signaling, which is closely related to AD pathology, can be influenced by PIK3CB through its interaction with the PI3K-AKT pathway and BCL-2, a protein known for its anti-apoptotic effects<sup>114</sup>. The intersection of these pathways highlights the multifaceted role of PIK3CB in AD, where its downregulation may contribute to the disease by impairing cell survival mechanisms and promoting neurodegeneration. |
| **VAV3** | Astrocytes, which are essential for neuronal support, glutamate uptake, and modulation of neuronal activity, show altered behavior in the absence of VAV3. Specifically, VAV3-deficient astrocytes exhibit enhanced regenerative capabilities and an altered cytokine release profile, including a complete lack of CXCL11, reduced levels of IL-6, and increased release of CCL5<sup>115</sup>. These changes can significantly influence the neuronal environment and potentially affect the progression of neurodegenerative diseases like AD. Given that VAV3 influences the release of cytokines and other signaling molecules from astrocytes, it is also plausible that VAV3 indirectly affects the inflammatory and immune responses in the AD brain, potentially modulating the formation and clearance of Aβ plaques<sup>116</sup>. Additionally, the altered axonal and dendritic morphology observed in VAV3-deficient neurons, such as increased axonal lengths and complexities, could impact neuronal connectivity and function, further contributing to the pathological mechanisms of AD<sup>115</sup>. Therefore, while direct evidence linking VAV3 to AD pathology is limited, its regulatory role in astrocyte function and neuron-glia interactions suggests that VAV3 could be an important modulator in the disease's progression, influencing both neuroinflammatory responses and neuronal structural integrity. |
| **GNG4** | GNG4 expression decreases in the brain with age and is notably downregulated in rodent models of AD<sup>117</sup>. This suggests a potential role in neurodegenerative processes, as GNG4 is involved in hemostasis and glucagon response, which are critical for cellular homeostasis. GNG4 expression is highest in the human hippocampus, a region crucial for memory and cognitive function and severely affected in AD<sup>117</sup>. A gender-specific aspect of GNG4's influence on cognitive decline has been observed; its association with cognitive decline was replicated in an all-female cohort but not in an all-male cohort, suggesting a possible interaction with estrogen, a hormone known to modify dementia risk<sup>117</sup>. Additionally, a weighted gene co-expression network analysis, have identified GNG4 as one of synapse-associated genes that is dysregulated in AD, specifically downregulated in the dorsolateral prefrontal cortex<sup>118</sup>. |

### Signaling Pathways

The Apelin signaling pathway regulates apoptosis, autophagy, synaptic plasticity, and neuroinflammation. Apelin-13, a key member of the apelin family, has significant neuroprotective functions that help prevent AD by modulating these cellular processes. The Apelin/APJ system influences several signaling pathways, such as PI3K/Akt, MAPK, and PKA, which are essential for cell proliferation and protection from excitotoxicity^25^. Alterations in apelin expression are linked to inflammatory responses, oxidative stress, Ca2+ signaling, and apoptosis, all related to AD pathology^26^. The intersection of the Apelin signaling pathway with the WNT signaling pathway suggests a broader regulatory network influencing AD-associated pathologies^27^. The apelinergic system’s involvement in brain physiology, including its protective effects against neurological disorders, highlights its importance in maintaining cognitive function and preventing neurodegeneration^28^.

The relaxin signaling pathway influences neuroinflammation, neurovascular integrity, and cognitive functions. Relaxin, a peptide hormone, modulates brain functions such as arousal, stress responses, and social recognition, which are critical in neuropsychiatric disorders^29^. It inhibits aberrant myofibroblast differentiation and collagen deposition through the TGF-β1/Smad2 axis and stimulates matrix metalloproteinases via the RXFP1-pERK-nNOS-NO-cGMP-dependent pathway, mitigating neuroinflammation and fibrosis, key features of AD pathology^30^. Relaxin also enhances neurovascular health by stimulating cAMP production and activating the PI3K/PKCζ pathway, leading to increased VEGF expression^31^. The pathway’s interaction with the WNT signaling pathway, crucial for cell adhesion and differentiation, further underscores its potential impact on AD.

The oxytocin (Oxt) signaling pathway influences social behavior, neuroinflammation, and cognitive function. Oxt administration has been shown to reverse learning and memory impairments in AD models, suggesting its potential as a therapeutic target^32^. Key mechanisms include inhibiting microglial activation and reducing inflammatory cytokine levels by blocking the ERK/p38 MAPK and COX-2/iNOS NF-κB pathways, which prevents cognitive impairment and delays hippocampal atrophy^33^. Oxt also reduces brain inflammation and corrects memory deficits by promoting Aβ deposition in dense core plaques, offering neuroprotective effects^34^. Chronic intranasal Oxt administration restores cognitive functions, reduces acetylcholinesterase activity, and lowers levels of β-amyloid and Tau proteins^35^. These effects are supported by decreased hippocampal ERK1/2 and GSK3β levels, reduced neuronal death, low caspase-3 activity, and improved histopathological profiles, highlighting Oxt’s potential in modulating AD pathology^35^.

Calcium signaling regulates neuronal function and survival. Dysregulation of calcium homeostasis is evident at all stages of AD and is linked to mitochondrial failure, oxidative stress, chronic neuroinflammation, and the formation of NFTs and Aβ plaques^36^. Glutamatergic NMDA receptor (NMDAR) activity is particularly significant, as NMDAR-mediated neurotoxicity is a key factor in AD progression^36^. Calcium dyshomeostasis is also associated with tau hyperphosphorylation, abnormal synaptic plasticity, and apoptosis^37^. Disruptions at the ER-mitochondria membrane contact site and decreased calcium-binding buffers further contribute to cellular toxicity in AD^38^.

### Cellular Processes

Circadian entrainment influences various physiological and pathological processes. Disruptions in circadian rhythms are common in AD, often preceding cognitive symptoms and exacerbating pathology through increased Aβ production, impaired clearance, and neuroinflammation^39^. Core circadian clock genes like BMAL1, PER, and CRY show altered expression in AD, contributing to symptoms such as disrupted sleep patterns, activity changes, and mood fluctuations^40^. AD model mice display novel circadian behaviors, including heightened sensitivity to light cues and faster re-entrainment to shifted light-dark cycles, indicating that AD pathology affects retinal light sensing^39^. Brain-wide spatial transcriptomics reveal progressive disruptions in diurnal transcriptional rhythms in AD, linking these alterations to disease pathology^41^. These findings suggest that targeting circadian clock genes and regulatory pathways could offer therapeutic strategies, such as optimizing drug administration timing or employing chronotherapeutics to mitigate disease progression and improve quality of life for AD patients.

Morphine addiction can significantly impact the development and progression of AD as opioids like morphine interfere with insulin signaling pathways via crosstalk between the insulin receptor and the mu-opioid receptor, crucial for neuronal health^42^. Morphine also affects neurotransmitter regulation, involving acetylcholine, norepinephrine, GABA, glutamate, and serotonin, which are implicated in AD, contributing to cognitive impairment and neuroinflammation^43^. Morphine downregulates BACE-1 and upregulates BACE-2 expression, affecting Aβ production through a nitric oxide-dependent mechanism, potentially leading to chronic vasoconstriction, brain hypoperfusion, and neuronal death^44^. Individuals with opioid use disorder have a significantly higher risk of developing AD and dementia, especially in younger populations^45^. Machine learning models suggest that including data on AD drugs and cognitive scores improves AD progression prediction, indicating that managing opioid addiction could help mitigate the disease’s advancement^46^.

Gastric acid secretion influences AD through its impact on gut health and the brain-gut axis. Proper gastric acid levels are essential for nutrient absorption, gut homeostasis, and protection against pathogens. Disruption in gastric acid secretion can lead to gut dysbiosis and increased gut permeability, which are linked to AD. For example, conditions like Helicobacter pylori infection, which alter gastric acid levels, can cause gut inflammation and subsequent neuroinflammation^47^. Gut inflammation in the gut can activate C/EBPβ/δ-secretase signaling, leading to the formation of Aβ and tau fibrils, which can then propagate to the brain via the vagus nerve, exacerbating AD^48^. Altered gastric acid secretion also affects gastrointestinal mucus production, compromising the gut barrier and increasing susceptibility to systemic inflammation^49^. Not to mention, the gut microbiota, influenced by gastric acid levels, plays a role in neuroinflammation and the formation of AD-related brain plaques and NFTs^50^.

Inflammatory mediators significantly regulate TRP channels, which particularly TRPV1 and TRPC6, are involved in neuroinflammation and calcium homeostasis disruption^51,52^. TRPV1 modulates neuroinflammation by influencing the production of inflammatory mediators and oxidative stress responses^52^. Its activation can rescue microglial dysfunction and restore immune responses, including phagocytic activity and autophagy, through the AKT/mTOR pathway, reducing amyloid pathology and reversing memory deficits in AD models^51,53^. TRPC6 affects calcium signaling pathways, which are disrupted in AD^51^. The regulation of TRP channels by inflammatory mediators also helps maintain the integrity of the BBB and neurovascular coupling, both compromised in AD. TRP channels are activated by reactive oxygen species, linking oxidative stress to neurodegenerative disease progression^54^. This interplay highlights the role of TRP channels in modulating neuroinflammation and oxidative stress, offering promising avenues for AD treatment.

### Dysfunctions in Neurotransmitter System

The development and progression of AD are intricately linked to dysfunctions in neurotransmitter systems, including glutamatergic, cholinergic, GABAergic, and dopaminergic synapses. Glutamatergic synapses, essential for cognitive and behavioral functions, are significantly affected in AD. Dysregulated glutamatergic mechanisms contribute to cognitive impairments and disease progression through interactions with neuronal hyperactivity, Aβ, tau, and glial dysfunction^55^. Aβ disrupts glutamate receptors like NMDA and AMPA, leading to calcium dyshomeostasis and impaired synaptic plasticity, characterized by suppressed long-term potentiation and enhanced long-term depression^56^. Additionally, altered glucose metabolism affects glutamate levels, exacerbating synaptic dysfunction in AD^57^.

Cholinergic synapses are also critically involved, with cholinergic atrophy accelerating cognitive decline. The cholinergic hypothesis posits that deficits in cholinergic signaling lead to abnormal tau phosphorylation, neuroinflammation, and cell apoptosis^58^. The basal forebrain cholinergic innervation of cortical areas is particularly vulnerable, and cholinergic receptor regulation is a hallmark of AD progression^59^.

GABAergic synapses, responsible for inhibitory signaling, are disrupted in AD due to alterations in the GABA_A_ receptor system and perineuronal nets, leading to synaptic hyperactivity and abnormal brain oscillations, contributing to cognitive deficits^60^. Although less studied, dopaminergic synapses also play a role in AD, with D2 dopaminergic receptors implicated in symptomatology^59^.

Synaptic dysfunction is a common pathogenic trait in AD, with synapse loss closely correlating with cognitive decline. The interplay between Aβ and tau at the synapse exacerbates synaptic deficits, making targeting these dysfunctions crucial for developing therapeutic strategies. Aberrant neurotransmission, including cholinergic, adrenergic, and glutamatergic networks, underpins cognitive decline in AD, with NMDAR dysfunction being particularly significant. Together, these neurotransmitter system alterations highlight the complexity of AD pathogenesis and the need for targeted interventions to mitigate synaptic dysfunction and cognitive decline.

### Viral Infections

Research indicates that HCMV may contribute to poorer cognitive abilities and augment tauopathy by interacting with TRA CDR3 and tau peptides^61^. Additionally, HCMV, along with other herpesviruses, has been shown to impact AD-related processes such as Aβ formation, neuronal death, and autophagy through virus-host protein-protein interactions^62^. Persistent HCMV infections can lead to the generation of AD hallmarks, including Aβ plaques and NFTs composed of hyperphosphorylated tau proteins, by exploiting pathways involved in oxidative stress and neuroinflammation^63^.

Kaposi Sarcoma-associated Herpesvirus (KSHV), a member of the Herpesviridae family, is known to impact AD-related processes such as Aβ formation, neuronal death, and autophagy, which are critical in the pathogenesis of AD^62^. The “infectious hypothesis” of AD suggests that pathogens, including viruses like KSHV, may act as seeds for Aβ aggregation, leading to plaque formation and cognitive decline^64^. Viral infections, including those caused by herpesviruses, can trigger neuroinflammatory pathways, disrupt the BBB, and activate microglia, leading to neural cell death and neurodegeneration^65^. Specifically, KSHV, along with other herpesviruses, has been shown to influence processes crucial for cellular homeostasis and dysfunction, potentially exacerbating AD pathology through virus-host protein-protein interactions^62^. Additionally, the reactivation of herpesviruses during acute infections, such as SARS-CoV-2, can create a synergistic pathogenic effect, further promoting neurodegenerative processes like Aβ formation and oxidative stress response^62^.

### Innate Immune Signaling Pathway

The chemokine signaling pathway plays a critical role in AD pathogenesis by driving neuroinflammation and regulating immune cell activity. Dysregulated chemokines, such as CCL5, CXCL1, and CXCL16, are found in both brain tissues and blood of AD patients, correlating with Aβ and tau pathology, and suggesting their potential as biomarkers for AD^66^. The CCL5/CCR5 axis is particularly notable for its dual role in normal physiology and neurodegeneration^67^. Chronic microglial activation, fueled by persistent Aβ deposition, leads to a loss of neuroprotective functions and increased neuronal damage^68^. Chemokines like CX3CL1 are vital in balancing microglial activity between neuroprotection and neurotoxicity^68^. Overexpression of chemokines can disrupt the BBB, facilitating immune cell infiltration and prolonged inflammation, which in turn enhances Aβ production, aggregation, impairs its clearance, and promotes tau hyperphosphorylation, contributing to neuronal loss and AD progression^69^. Elevated levels of chemokines in AD patient plasma further underscore their role in the disease^70^.

### Hemostasis

Platelets are a major peripheral source of Aβ, providing about 90% of circulating Aβ, which is a hallmark of AD^71,72^. Elevated platelet activity, particularly in APOE4 carriers, correlates with disease severity and cognitive decline, making platelet activity a potential marker for AD progression^71^. Platelets show altered levels of amyloid precursor protein, metabolic enzymes, oxidative stress markers, and neurotransmitters, reflecting changes seen in the central nervous system of AD patients^73^. The PI3K/AKT pathway, which regulates platelet activity, influences Aβ production by regulating APP, BACE-1, ADAMs, and γ-secretase. ROS-induced oxidative stress, a key factor in AD, also leads to platelet hyperactivity, worsening neuroinflammation and neurodegeneration^72,74^. Additionally, in conditions like type 2 diabetes mellitus, abnormal platelet reactivity and insulin resistance contribute to vascular dysfunction and Aβ aggregation, accelerating AD progression^75^.

### Model validation: gene validation

The identification of the top 20 genes associated with AD in females and males, listed in **Table 8**, represents a significant step forward in understanding the genetic underpinnings of this condition. The genes were ranked using our novel graph AI model, which integrated multi-omic data to highlight key biomarkers and signaling interactions relevant to AD. Notably, the same genes were identified for both sexes, underscoring their critical role in AD pathogenesis. **Table 8** provides a comprehensive overview of these genes, detailing their functions and specific relations to AD. The function of each gene and its contribution to AD pathology are pivotal for elucidating the complex biological mechanisms driving the disease. The consistency of gene rankings between females and males emphasizes the universal importance of these biomarkers in AD. This uniformity suggests that the identified genes play fundamental roles in the disease’s progression, regardless of sex. This finding enhances the potential for developing targeted therapies that could benefit a broad patient population.

## Declarations

### Ethics approval and consent to participate

Not applicable, as no patient data was used in this research. Cell line used in this study is not relevant material under the Human Tissue Act, so no ethical approval was required.

### Consent for publication

Not applicable, as no patient data were used in this research.

### Availability of data and material

Check the Table 1 for details in the Methodology and Materials section.

### Funding

This study was partially supported by NIA R56AG065352 (to Li),

1R21AG078799-01A1 (to Li/Province), NINDS 1RM1NS132962-01 (to

Dickson/Marco/Cooper/Li).

## Appendix

### Section A

**Figure S1.**
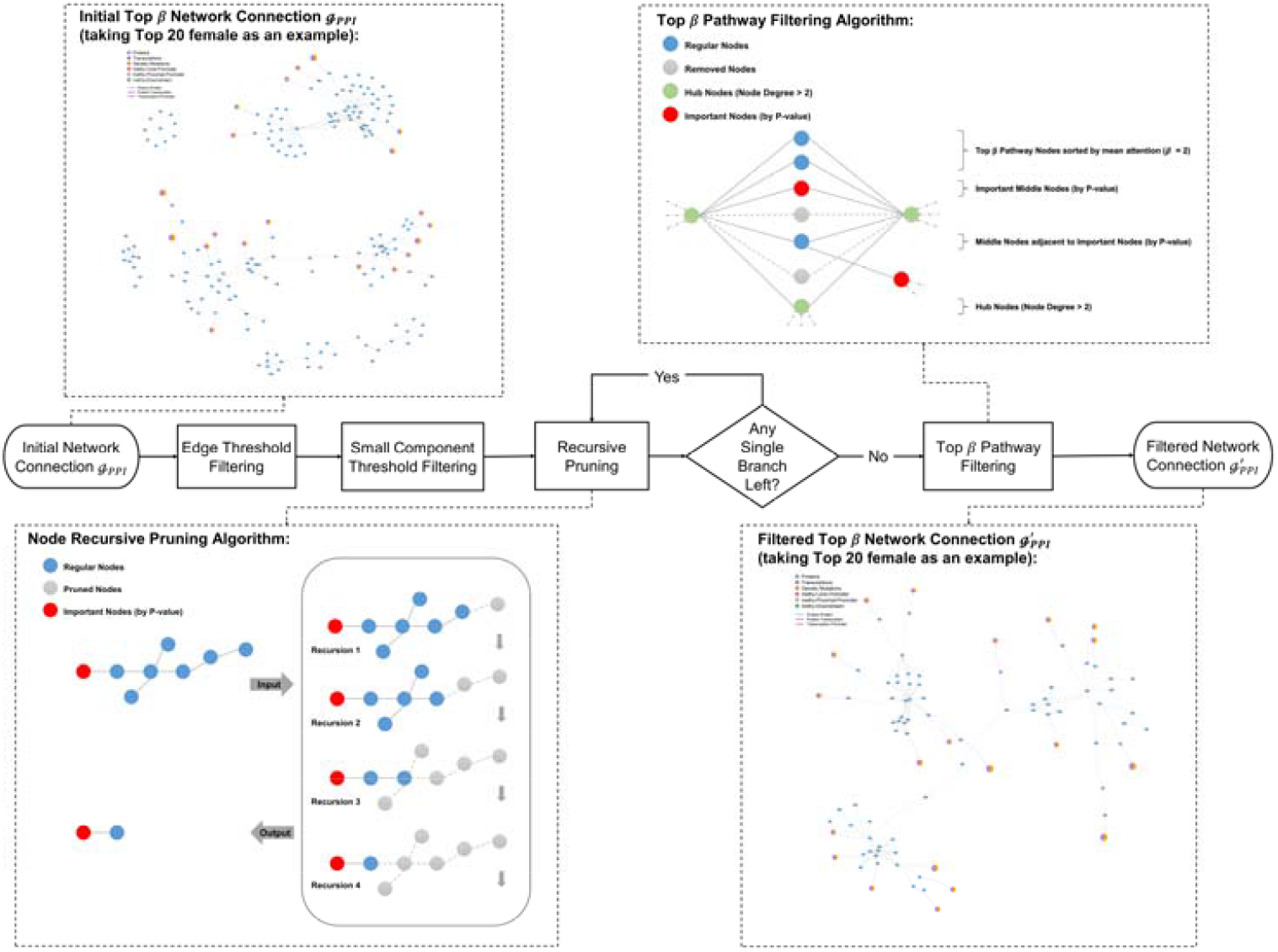
Diagram of the processing procedures for core signaling networks visualization.

Figure S1 illustrates the process of filtering and pruning network connections. Initially, the network connections are processed through edge threshold filtering and small component threshold filtering. Then, a recursive node pruning algorithm is applied to remove insignificant nodes. Next, it checks if any single branch remains; if so, pathway filtering is performed. Finally, a filtered network connection is obtained. The diagram also provides examples of the initial and filtered network connections, along with detailed steps of the recursive node pruning and pathway filtering algorithms.

### Section B

**Table S1.**
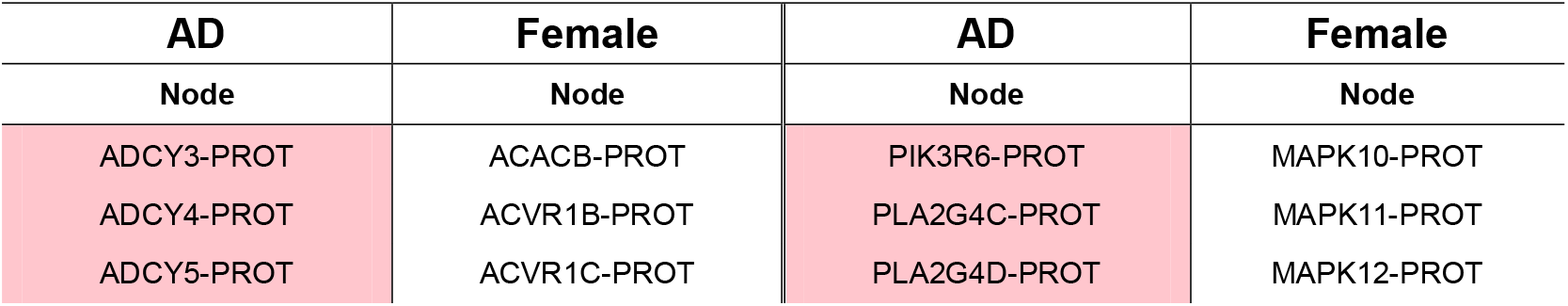
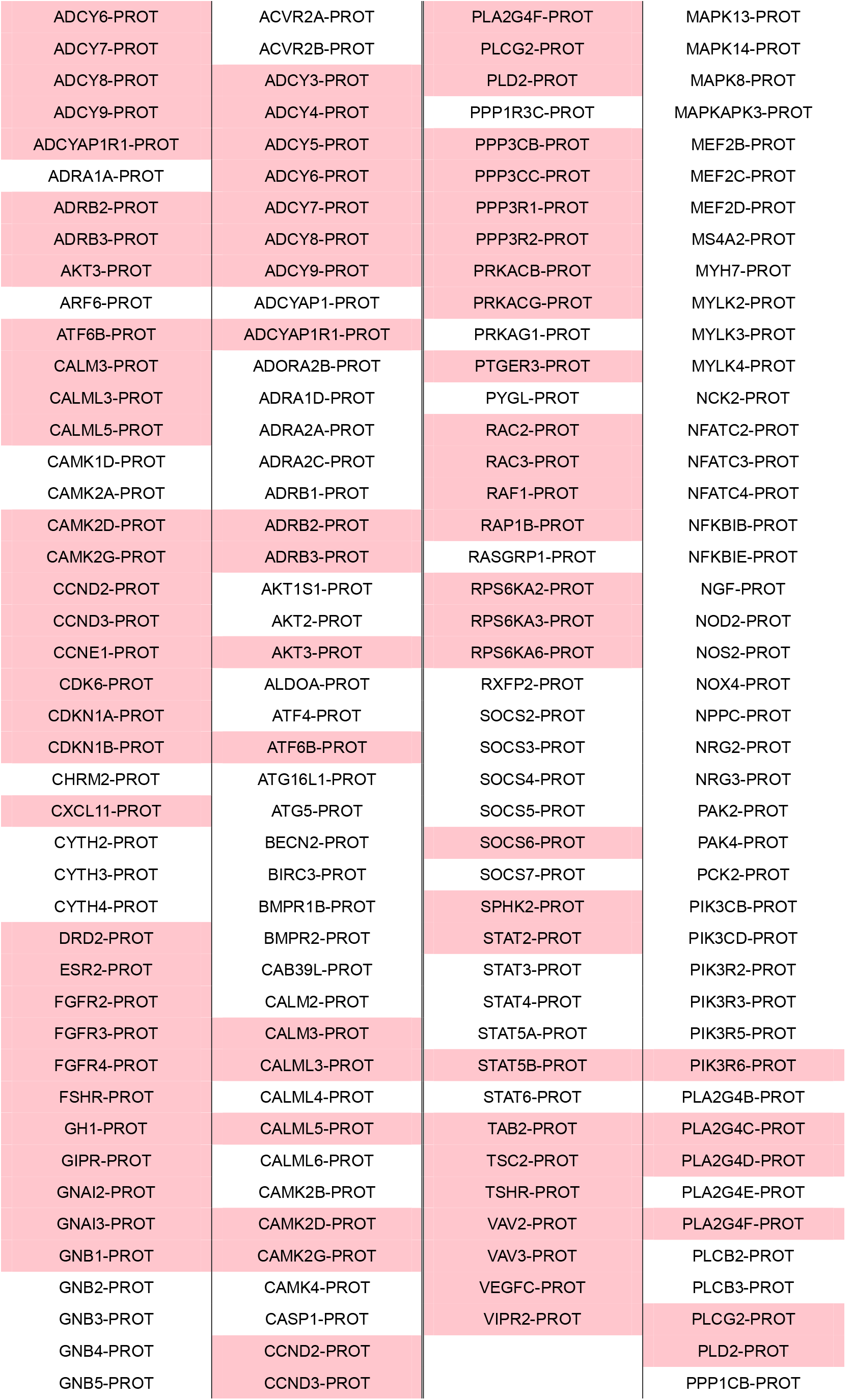
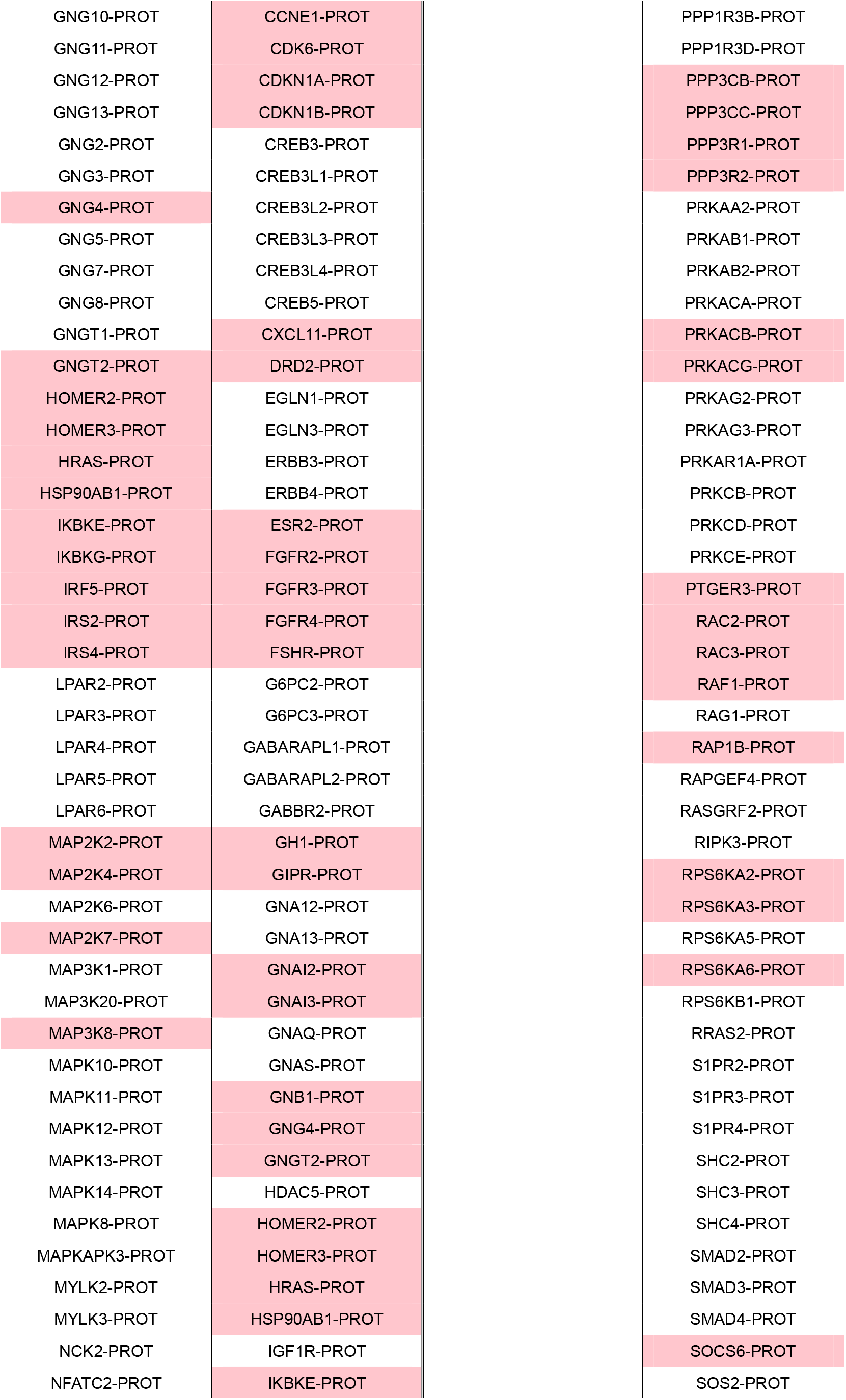
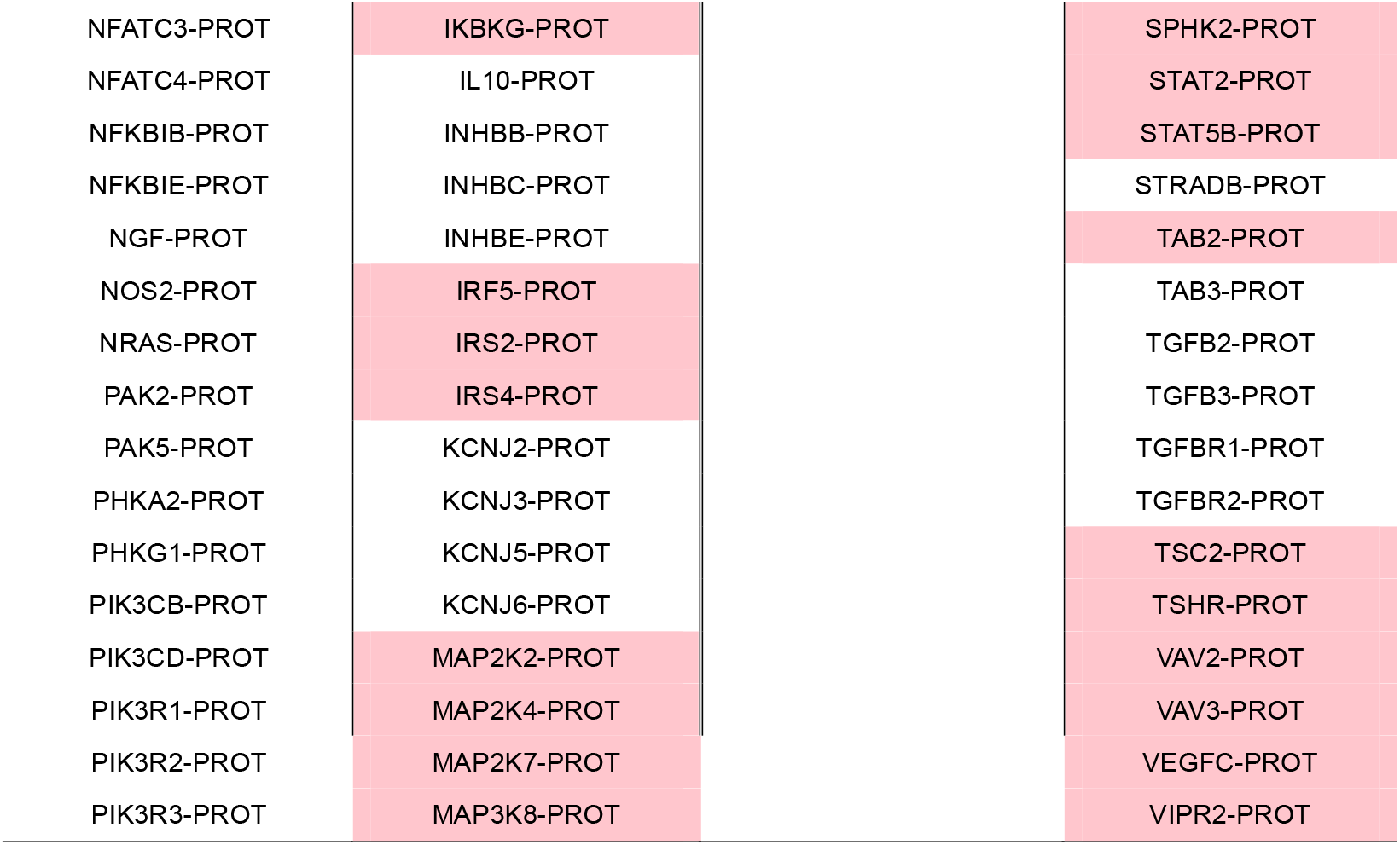
Top 70 AD-Female overlapping protein nodes (overlapped gene features marked with pink)

**Table S2.**
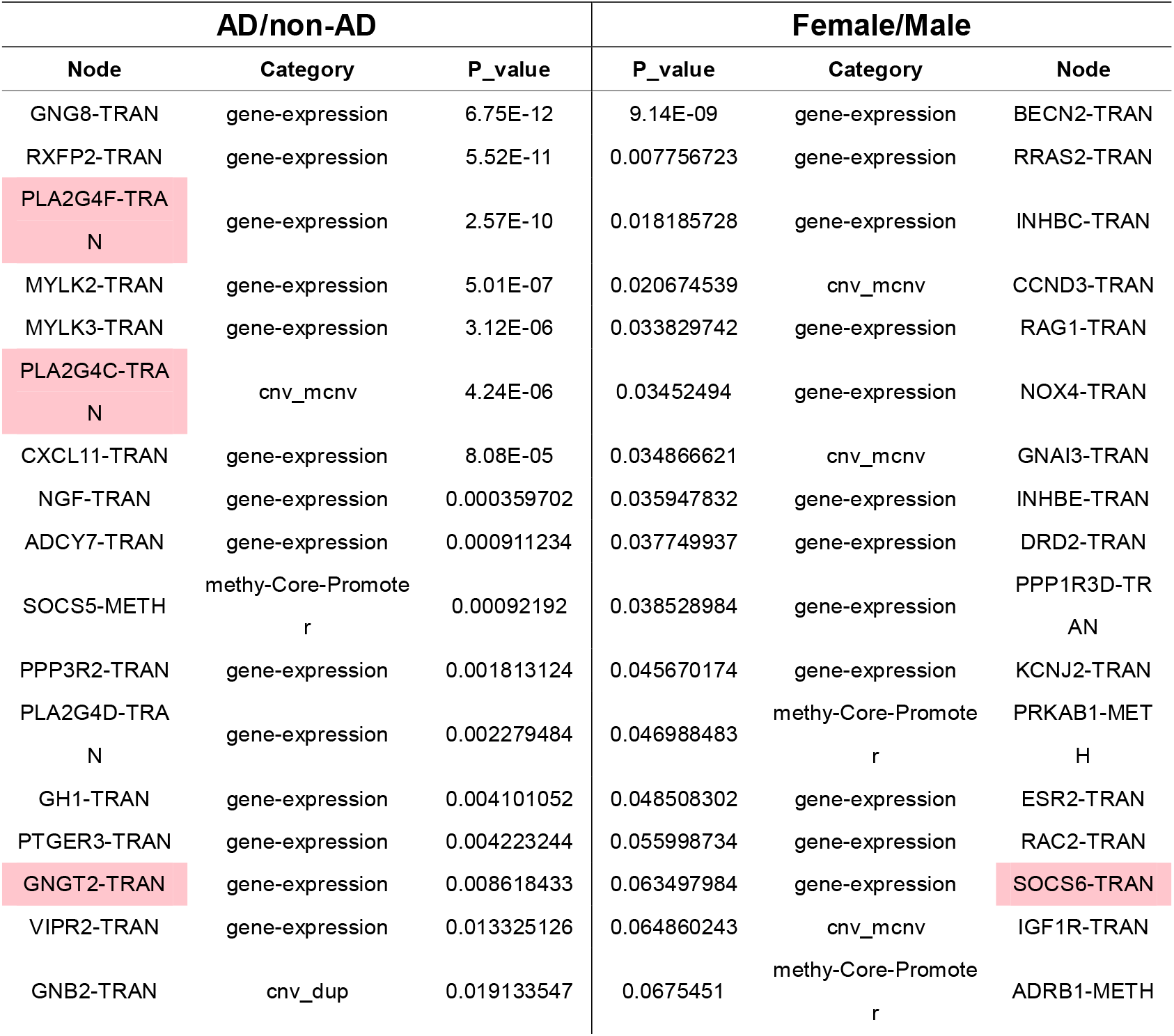
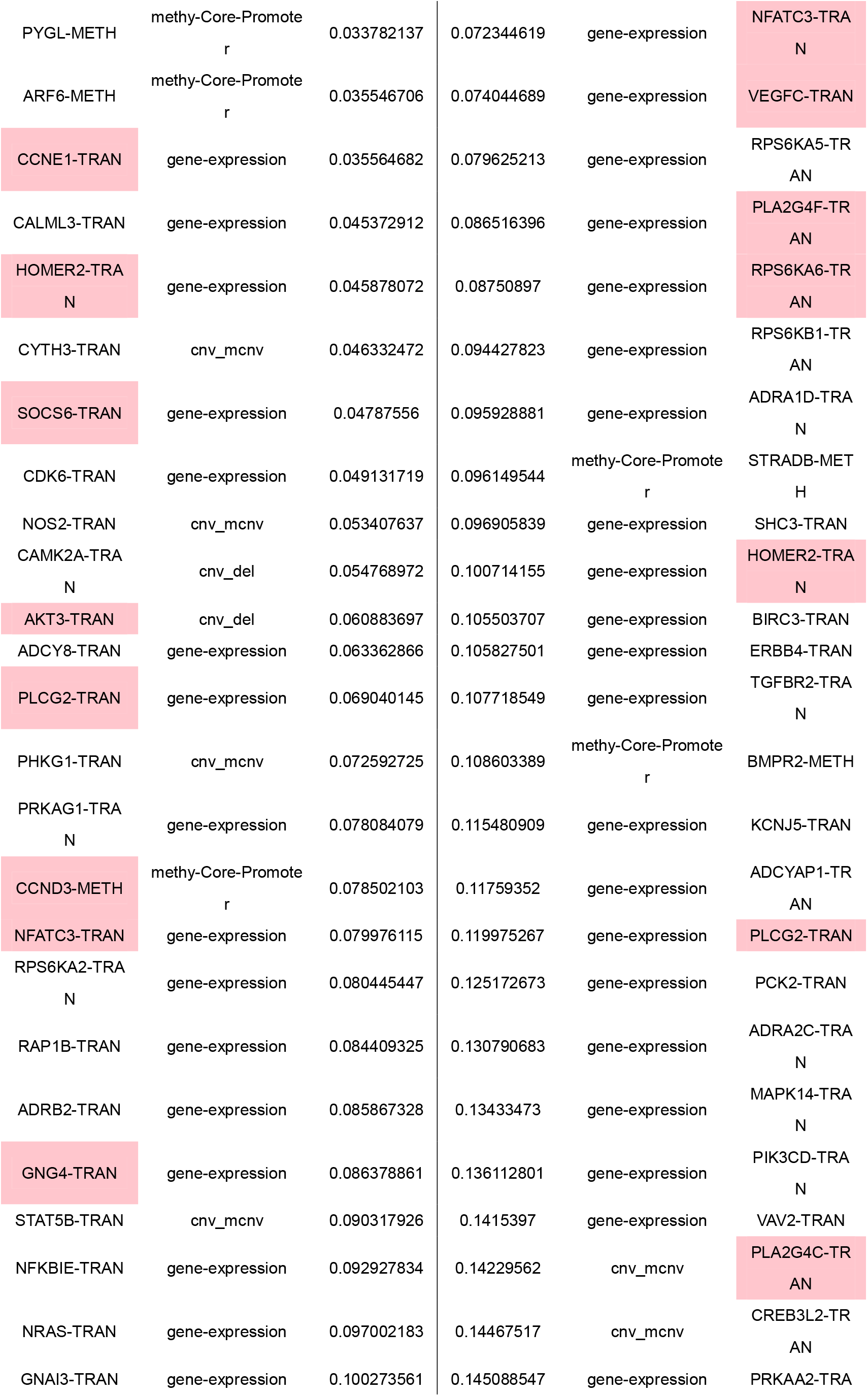
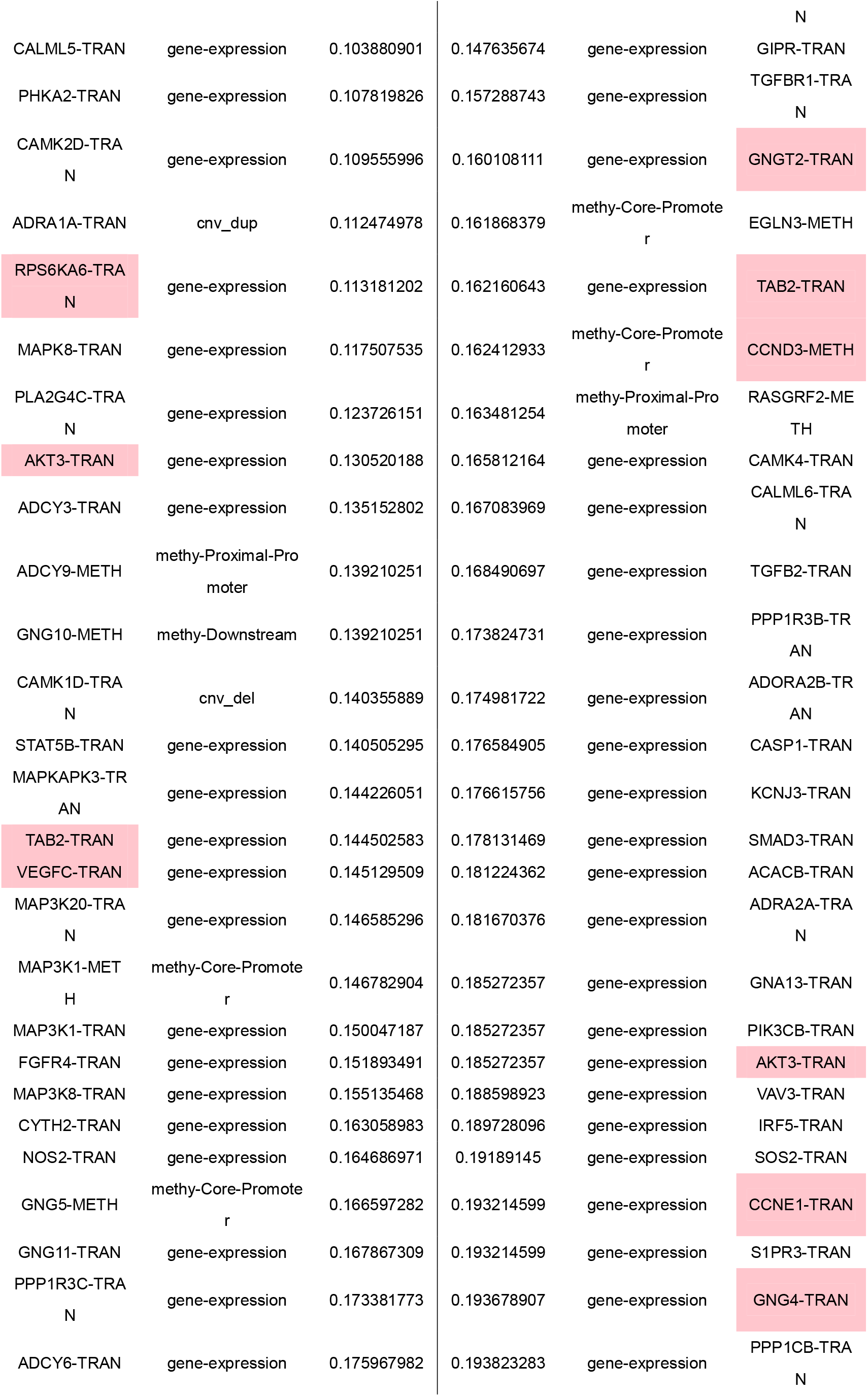
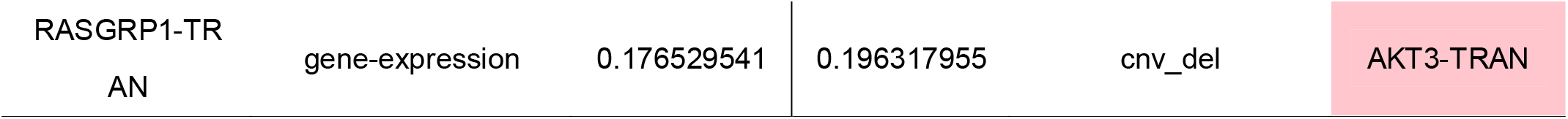
Top 70 AD/Non-AD and female/male overlapped gene features (overlapped gene features marked with pink)

### Section C

**Table S3.** Pathway enrichment analysis results using ShinyGO 0.80 and KEGG database

| Type | Pathway | Number of Genes | Genes | P-Value | FDR |
| --- | --- | --- | --- | --- | --- |
| Signaling Pathways | hsa05200: Pathways in cancer | 28 | GNG11, ADCY3, CCNE1, CDK6, ADCY8, FGFR4, STAT5B, ADCY9, GNG8, GNB2, RASGRP1, ADCY6, CALML3, NOS2, CAMK2D, PTGER3, NRAS, GNAI3, GNG5, CCND3, GNG10, CALML5, ADCY7, MAPK8, CAMK2A, GNGT2, VEGFC, PLCG2 | 3.36E-27 | 1.13E-24 |
| Signaling Pathways | hsa04921: Oxytocin signaling pathway | 20 | ADCY3, ADCY8, PLA2G4D, ADCY9, NFATC3, ADCY6, CALML3, PRKAG1, CAMK2D, NRAS, GNAI3, PPP3R2, MYLK2, CALML5, ADCY7, MYLK3, PLA2G4F, CAMK2A, PLA2G4C, CAMK1D | 1.55E-26 | 2.6E-24 |
| Signaling Pathways | hsa04014: Ras signaling pathway | 20 | GNG11, RAP1B, PLA2G4D, FGFR4, ARF6, GNG8, GNB2, RASGRP1, CALML3, NGF, NRAS, GNG5, GNG10, CALML5, MAPK8, PLA2G4F, GNGT2, PLA2G4C, VEGFC, PLCG2 | 3.5E-23 | 2.35E-21 |
| Signaling Pathways | hsa04371: Apelin signaling pathway | 19 | GNG11, ADCY3, ADCY8, ADCY9, GNG8, GNB2, ADCY6, CALML3, PRKAG1, NOS2, NRAS, GNAI3, GNG5, GNG10, MYLK2, CALML5, ADCY7, MYLK3, GNGT2 | 8.58E-26 | 9.61E-24 |
| Viral Infections | hsa05163: Human cytomegalovirus infection | 19 | GNG11, ADCY3, CDK6, ADCY8, ADCY9, NFATC3, GNG8, GNB2, ADCY6, CALML3, PTGER3, NRAS, GNAI3, GNG5, GNG10, PPP3R2, CALML5, ADCY7, GNGT2 | 6.63E-22 | 2.79E-20 |
| Signaling Pathways | hsa04010: MAPK signaling pathway | 19 | RAP1B, RPS6KA6, MAP3K8, PLA2G4D, FGFR4, NFATC3, RASGRP1, TAB2, NGF, NRAS, PPP3R2, ZAK, MAPK8, PLA2G4F, MAP3K1, MAPKAPK3, RPS6KA2, PLA2G4C, VEGFC | 9.26E-20 | 2.22E-18 |
| Signaling Pathways | hsa04020: Calcium signaling pathway | 18 | ADCY3, ADCY8, ADCY9, ADRB2, CALML3, NOS2, CAMK2D, PTGER3, PPP3R2, MYLK2, PHKA2, CALML5, ADRA1A, ADCY7, MYLK3, CAMK2A, CAMK1D, PLCG2 | 2.98E-21 | 1E-19 |
| Cellular Processes | hsa04724: Glutamatergic synapse | 17 | GNG11, ADCY3, ADCY8, PLA2G4D, ADCY9, GNG8, GNB2, HOMER2, ADCY6, GNAI3, GNG5, GNG10, PPP3R2, ADCY7, PLA2G4F, GNGT2, PLA2G4C | 1.83E-23 | 1.54E-21 |
| Signaling Pathways | hsa04926: Relaxin signaling pathway | 17 | GNG11, ADCY3, ADCY8, ADCY9, RXFP2, GNG8, GNB2, ADCY6, NOS2, NRAS, GNAI3, GNG5, GNG10, ADCY7, MAPK8, GNGT2, VEGFC | 1.68E-22 | 8.06E-21 |
| Innate Immune Signaling Pathway | hsa04062: Chemokine signaling pathway | 17 | GNG11, RAP1B, ADCY3, ADCY8, STAT5B, ADCY9, GNG8, GNB2, CXCL11, ADCY6, NRAS, GNAI3, GNG5, GNG10, ADCY7, GNGT2, PLCG2 | 5.99E-20 | 1.55E-18 |
| Cellular Processes | hsa04713: Circadian entrainment | 16 | GNG11, ADCY3, ADCY8, ADCY9, GNG8, GNB2, ADCY6, CALML3, CAMK2D, GNAI3, GNG5, GNG10, CALML5, ADCY7, CAMK2A, GNGT2 | 6.56E-23 | 3.68E-21 |
| Signaling Pathways | hsa04912: GnRH signaling pathway | 15 | ADCY3, ADCY8, PLA2G4D, ADCY9, ADCY6, CALML3, CAMK2D, NRAS, CALML5, ADCY7, MAPK8, PLA2G4F, CAMK2A, MAP3K1, PLA2G4C | 2.52E-21 | 9.42E-20 |
| Innate Immune Signaling Pathway | hsa04750: Inflammatory mediator regulation of TRP channels | 15 | ADCY3, ADCY8, PLA2G4D, ADCY9, ADCY6, CALML3, CAMK2D, NGF, CALML5, ADCY7, MAPK8, PLA2G4F, CAMK2A, PLA2G4C, PLCG2 | 5.31E-21 | 1.62E-19 |
| Cellular Processes | hsa04725: Cholinergic synapse | 15 | GNG11, ADCY3, ADCY8, ADCY9, GNG8, GNB2, ADCY6, CAMK2D, NRAS, GNAI3, GNG5, GNG10, ADCY7, CAMK2A, GNGT2 | 4.57E-20 | 1.28E-18 |
| Viral Infections | hsa05170: Human immunodeficiency virus 1 infection | 15 | GNG11, NFATC3, GNG8, GNB2, CALML3, TAB2, NRAS, GNAI3, GNG5, GNG10, PPP3R2, CALML5, MAPK8, GNGT2, PLCG2 | 2.43E-16 | 4.53E-15 |
| Hemostasis | hsa04611: Platelet activation | 14 | RAP1B, ADCY3, ADCY8, PLA2G4D, ADCY9, RASGRP1, ADCY6, GNAI3, MYLK2, ADCY7, MYLK3, PLA2G4F, PLA2G4C, PLCG2 | 8.3E-18 | 1.86E-16 |
| Signaling Pathways | hsa04722: Neurotrophin | 13 | RAP1B, RPS6KA6, NFKBIE, CALML3, CAMK2D, NGF, NRAS, CALML5, MAPK8, CAMK2A, MAP3K1, RPS6KA2, PLCG2 | 1.48E-16 | 2.92E-15 |

## References

1. Zeliger, H. Oxidative Stress: Its Mechanisms and Impacts on Human Health and Disease Onset. (Academic Press, 2022).

2. MAPPING A BETTER FUTURE FOR DEMENTIA CARE NAVIGATION.

3. Kipf, T. N. & Welling, M. Semi-supervised classification with graph convolutional networks. 5th International Conference on Learning Representations, ICLR 2017 - Conference Track Proceedings 1–14 (2017).

4. Hamilton, W. L., Ying, R. & Leskovec, J. Inductive representation learning on large graphs. Adv Neural Inf Process Syst 2017-Decem, 1025–1035 (2017).

5. Veličković, P. et al. Graph attention networks. 6th International Conference on Learning Representations, ICLR 2018 - Conference Track Proceedings 1–12 (2018).

6. Xu, K., Hu, W., Leskovec, J. & Jegelka, S. HOW POWERFUL ARE GRAPH NEURAL NETWORKS?

7. Wang, T. et al. MOGONET integrates multi-omics data using graph convolutional networks allowing patient classification and biomarker identification. Nat Commun 12, 3445 (2021).

8. Li, X. et al. MoGCN: a multi-omics integration method based on graph convolutional network for cancer subtype analysis. Front Genet 13, 806842 (2022).

9. Gao, H. et al. A universal framework for single-cell multi-omics data integration with graph convolutional networks. Brief Bioinform 24, bbad081 (2023).

10. Rajadhyaksha, N. & Chitkara, A. Graph Contrastive Learning for Multi-omics Data. arXiv preprint 2301.02242 (2023).

11. Li, G., Muller, M., Thabet, A. & Ghanem, B. DeepGCNs: Can GCNs go as deep as CNNs? Proceedings of the IEEE International Conference on Computer Vision 2019-Octob, 9266–9275 (2019).

12. Cai, C. & Wang, Y. A Note on Over-Smoothing for Graph Neural Networks. (2020).

13. Abu-El-Haija, S. et al. Mixhop: Higher-order graph convolutional architectures via sparsified neighborhood mixing. in international conference on machine learning 21–29 (PMLR, 2019).

14. Morris, C. et al. Weisfeiler and Leman Go Neural: Higher-Order Graph Neural Networks. www.aaai.org.

15. Chien, E., Peng, J., Li, P. & Milenkovic, O. Adaptive universal generalized pagerank graph neural network. arXiv preprint 2006.07988 (2020).

16. Nikolentzos, G., Dasoulas, G. & Vazirgiannis, M. k-hop graph neural networks. Neural Networks 130, 195–205 (2020).

17. Brossard, R., Frigo, O. & Dehaene, D. Graph convolutions that can finally model local structure. arXiv preprint 2011.15069 (2020).

18. Wang, G., Ying, R., Huang, J. & Leskovec, J. Multi-hop attention graph neural network. arXiv preprint 2009.14332 (2020).

19. Feng, J., Chen, Y., Li, F., Sarkar, A. & Zhang, M. How Powerful Are K-Hop Message Passing Graph Neural Networks.

20. Zhang, H. et al. mosGraphGen: a novel tool to generate multi-omic signaling graphs to facilitate integrative and interpretable graph AI model development. doi:10.1101/2024.05.15.594360.

21. Zhang, H. et al. M3NetFlow: a novel multi-scale multi-hop multi-omics graph AI model for omics data integration and interpretation. doi:10.1101/2023.06.15.545130.

22. Zhang, H., Chen, Y., Payne, P. & Li, F. Mining signaling flow to interpret mechanisms of synergy of drug combinations using deep graph neural networks. doi:10.1101/2021.03.25.437003.

23. Kipf, T. N. & Welling, M. Semi-supervised classification with graph convolutional networks. arXiv preprint 1609.02907 (2016).

24. Shi, Y. et al. Masked Label Prediction: Unified Message Passing Model for Semi-Supervised Classification.

25. Kinjo, T., Higashi, H., Uno, K. & Kuramoto, N. Apelin/Apelin Receptor System: Molecular Characteristics, Physiological Roles, and Prospects as a Target for Disease Prevention and Pharmacotherapy. Curr Mol Pharmacol 14, 210–219 (2021).

26. Luo, H., Han, L. & Xu, J. Apelin/APJ system: A novel promising target for neurodegenerative diseases. Journal of Cellular Physiology vol. 235 638–657 Preprint at 10.1002/jcp.29001 (2020).

27. Kostes, W. W. & Brafman, D. A. The Multifaceted Role of WNT Signaling in Alzheimer’s Disease Onset and Age-Related Progression. Cells vol. 12 Preprint at 10.3390/cells12081204 (2023).

28. Ivanov, M. N., Stoyanov, D. S., Pavlov, S. P. & Tonchev, Anton. B. Distribution, Function, and Expression of the Apelinergic System in the Healthy and Diseased Mammalian Brain. Genes (Basel) 13, 2172 (2022).

29. Blasiak, A. et al. Relaxin ligand/receptor systems in the developing teleost fish brain: Conserved features with mammals and a platform to address neuropeptide system functions. Frontiers in Molecular Neuroscience vol. 15 Preprint at 10.3389/fnmol.2022.984524 (2022).

30. Chow, B. S. M. et al. Relaxin signals through a RXFP1-pERK-nNOS-NO-cGMP-dependent pathway to up-regulate matrix metalloproteinases: The additional involvement of iNOS. PLoS One 7, (2012).

31. Dessauer, C. W. & Nguyen, B. T. Relaxin stimulates multiple signaling pathways: Activation of cAMP, PI3K, and PKCζ in THP-1 cells. in Annals of the New York Academy of Sciences vol. 1041 272–279 (New York Academy of Sciences, 2005).

32. Takahashi, J., Yamada, D., Nagano, W. & Saitoh, A. The Role of Oxytocin in Alzheimer’s Disease and Its Relationship with Social Interaction. Cells vol. 12 Preprint at 10.3390/cells12202426 (2023).

33. Ye, C. et al. Oxytocin Nanogels Inhibit Innate Inflammatory Response for Early Intervention in Alzheimer’s Disease. ACS Appl Mater Interfaces 14, 21822–21835 (2022).

34. Clara Selles, M. et al. Oxytocin attenuates microglial activation and restores social and non-social memory in the APP/PS1 mouse model of Alzheimer’s disease. doi:10.1101/2022.05.07.491031.

35. El-Ganainy, S. O. et al. Intranasal Oxytocin Attenuates Cognitive Impairment, β-Amyloid Burden and Tau Deposition in Female Rats with Alzheimer’s Disease: Interplay of ERK1/2/GSK3β/Caspase-3. Neurochem Res 47, 2345–2356 (2022).

36. Baracaldo-Santamaría, D. et al. Role of Calcium Modulation in the Pathophysiology and Treatment of Alzheimer’s Disease. International Journal of Molecular Sciences vol. 24 Preprint at 10.3390/ijms24109067 (2023).

37. Ge, M. et al. Role of Calcium Homeostasis in Alzheimer’s Disease. Neuropsychiatr Dis Treat 18, 487–498 (2022).

38. Huang, D.-X. et al. Calcium Signaling Regulated by Cellular Membrane Systems and Calcium Homeostasis Perturbed in Alzheimer’s Disease. Front Cell Dev Biol (2022) doi:10.3389/fcell.2022.834962.

39. Weigel, T. K., Guo, C. L., Güler, A. D. & Ferris, H. A. Altered circadian behavior and light sensing in mouse models of Alzheimer’s disease. Front Aging Neurosci 15, (2023).

40. Aili, A. & Zeng, Z. Circadian Clock Gene Dysregulation in Alzheimer’s Disease: Insights and Implications. (2024).

41. Romero, H., Gerber, A., Akhmetova, L., Mukamel, E. & Desplats, P. Spatial transcriptomics identifies disrupted circadian gene expression in a mouse model of Alzheimer’s disease. Alzheimer’s & Dementia 19, (2023).

42. Salarinasab, S. et al. Interaction of opioid with insulin/IGFs signaling in Alzheimer’s disease. Journal of Molecular Neuroscience vol. 70 819–834 Preprint at 10.1007/s12031-020-01478-y (2020).

43. Cai, Z. & Ratka, A. Opioid system and Alzheimer’s disease. NeuroMolecular Medicine vol. 14 91–111 Preprint at 10.1007/s12017-012-8180-3 (2012).

44. Pakabcdef, T., Cadetadef, P., Mantioneab, K. J. & Stefanoeg, G. B. Morphine via Nitric Oxide Modulates B-Amyloid Metabolism: A Novel Protective Mechanism for Alzheimer’s Disease. http://www.medscimonit.com/abstract/index/idArt/429256 (2005).

45. Qeadan, F. et al. Exploring the Association Between Opioid Use Disorder and Alzheimer’s Disease and Dementia Among a National Sample of the U.S. Population. Journal of Alzheimer’s Disease 96, 229–244 (2023).

46. El-Sappagh, S. et al. The Role of Medication Data to Enhance the Prediction of Alzheimer’s Progression Using Machine Learning. Comput Intell Neurosci 2021, (2021).

47. Schubert, M. L. Physiologic, pathophysiologic, and pharmacologic regulation of gastric acid secretion. Current Opinion in Gastroenterology vol. 33 430–438 Preprint at 10.1097/MOG.0000000000000392 (2017).

48. Chen, C. et al. Gut inflammation triggers C/EBPβ/δ-secretase-dependent gut-to-brain propagation of Aβ and Tau fibrils in Alzheimer’s disease. EMBO J 40, (2021).

49. Homolak, J. et al. Altered Secretion, Constitution, and Functional Properties of the Gastrointestinal Mucus in a Rat Model of Sporadic Alzheimer’s Disease. ACS Chem Neurosci 14, 2667–2682 (2023).

50. Bulgart, H. R., Neczypor, E. W., Wold, L. E. & Mackos, A. R. Microbial involvement in Alzheimer disease development and progression. Mol Neurodegener 15, 42 (2020).

51. Lu, R., He, Q. & Wang, J. TRPC Channels and Alzheimer’s Disease. Advances in Experimental Medicine and Biology 73–83 (2017) doi:10.1007/978-94-024-1088-4_7.

52. Duitama, M. et al. TRP Channels Role in Pain Associated With Neurodegenerative Diseases. Frontiers in Neuroscience vol. 14 reprint at 10.3389/fnins.2020.00782 (2020).

53. Lu, J., Zhou, W., Dou, F., Wang, C. & Yu, Z. TRPV1 sustains microglial metabolic reprogramming in Alzheimer’s disease. EMBO Rep 22, (2021).

54. Negri, S., Sanford, M., Shi, H. & Tarantini, S. The role of endothelial TRP channels in age-related vascular cognitive impairment and dementia. Frontiers in Aging Neuroscience vol. 15 reprint at 10.3389/fnagi.2023.1149820 (2023).

55. Pinky, P. D. et al. Recent Insights on Glutamatergic Dysfunction in Alzheimer’s Disease and Therapeutic Implications. Neuroscientist vol. 29 461–471 Preprint at 10.1177/10738584211069897 (2023).

56. Zhang, H. et al. Role of Aβ in Alzheimer’s-related synaptic dysfunction. Frontiers in Cell and Developmental Biology vol. 10 reprint at 10.3389/fcell.2022.964075 (2022).

57. Bukke, V. N. et al. The dual role of glutamatergic neurotransmission in Alzheimer’s disease: From pathophysiology to pharmacotherapy. International Journal of Molecular Sciences vol. 21 1–29 Preprint at 10.3390/ijms21207452 (2020).

58. Chen, Z. R., Huang, J. B., Yang, S. L. & Hong, F. F. Role of Cholinergic Signaling in Alzheimer’s Disease. Molecules vol. 27 reprint at 10.3390/molecules27061816 (2022).

59. Lombardero, L., Llorente-Ovejero, A., Manuel, I. & Rodríguez-Puertas, R. Chapter 28 - Neurotransmitter receptors in Alzheimer’s disease: from glutamatergic to cholinergic receptors. in Genetics, Neurology, Behavior, and Diet in Dementia (eds. Martin, C.R. & Preedy, V.R.) 441–456 (Academic Press, 2020). doi:10.1016/B978-0-12-815868-5.00028-1.

60. Ali, A. B., Islam, A. & Constanti, A. The fate of interneurons, GABAA receptor sub-types and perineuronal nets in Alzheimer’s disease. Brain Pathology vol. 33 Preprint at 10.1111/bpa.13129 (2023).

61. Blanck, G., Huda, T. I., Chobrutskiy, B. I. & Chobrutskiy, A. CMV as a factor in the development of Alzheimer’s disease? Med Hypotheses 178, (2023).

62. Onisiforou, A. & Zanos, P. From Viral Infections to Alzheimer’s Disease: Unveiling the Mechanistic Links Through Systems Bioinformatics. doi:10.1101/2023.12.05.570187.

63. Athanasiou, E., Gargalionis, A. N., Anastassopoulou, C., Tsakris, A. & Boufidou, F. New Insights into the Molecular Interplay between Human Herpesviruses and Alzheimer’s Disease—A Narrative Review. Brain Sciences vol. 12 Preprint at 10.3390/brainsci12081010 (2022).

64. Piotrowski, S. L., Tucker, A. & Jacobson, S. The elusive role of herpesviruses in Alzheimer’s disease: current evidence and future directions. NeuroImmune Pharmacology and Therapeutics 2, 253–266 (2023).

65. Du, C. Virus-induced Alzheimer’s disease: Potential roles of viral infections in AD neuropathogenesis from two aspects: Aberrant protein accumulations with neuroinflammatory response and virus-induced ablation of adult neurogenesis. AIP Conf Proc 2511, 020063 (2022).

66. Li, X. et al. Convergent transcriptomic and genomic evidence supporting a dysregulation of CXCL16 and CCL5 in Alzheimer’s disease. Alzheimers Res Ther 15, (2023).

67. Ma, W. et al. The intricate role of CCL5/CCR5 axis in Alzheimer disease. Journal of Neuropathology and Experimental Neurology vol. 82 894–900 Preprint at 10.1093/jnen/nlad071 (2023).

68. Bivona, G., Iemmolo, M. & Ghersi, G. CX3CL1 Pathway as a Molecular Target for Treatment Strategies in Alzheimer’s Disease. International Journal of Molecular Sciences vol. 24 Preprint at 10.3390/ijms24098230 (2023).

69. Wojcieszak, J., Kuczyńska, K. & B Zawilska, J. Role of Chemokines in the Development and Progression of Alzheimer’s Disease. Journal of Molecular Neuroscience 72, 1929–1951 (2022).

70. Wang, H., Zong, Y., Zhu, L., Wang, W. & Han, Y. Chemokines in patients with Alzheimer’s disease: A meta-analysis. Front Aging Neurosci 15, (2023).

71. Fu, J. et al. Correlation analysis of peripheral platelet markers and disease phenotypes in Alzheimer’s disease. Alzheimer’s and Dementia 20, 4366–4372 (2024).

72. Beura, S. K. et al. Redefining oxidative stress in Alzheimer’s disease: Targeting platelet reactive oxygen species for novel therapeutic options. Life Sciences vol. 306 Preprint at 10.1016/j.lfs.2022.120855 (2022).

73. Fu, J. et al. Meta-analysis and systematic review of peripheral platelet-associated biomarkers to explore the pathophysiology of alzheimer’s disease. BMC Neurol 23, (2023).

74. Khezri, M. R., Esmaeili, A. & Ghasemnejad-Berenji, M. Platelet Activation and Alzheimer’s Disease: The Probable Role of PI3K/AKT Pathway. Journal of Alzheimer’s Disease 90, 529–534 (2022).

75. Carbone, M. G., Pomara, N., Callegari, C., Marazziti, D. & Imbimbo, B. Pietro. TYPE 2 DIABETES MELLITUS, PLATELET ACTIVATION AND ALZHEIMER’S DISEASE: A POSSIBLE CONNECTION. Clin Neuropsychiatry 19, 370–378 (2022).

76. Hadar, A. et al. RGS2 expression predicts amyloid-β sensitivity, MCI and Alzheimer’s disease: Genome-wide transcriptomic profiling and bioinformatics data mining. Transl Psychiatry 6, (2016).

77. Yao, J., Chen, S. R. W., Yao, J. & Chen, S. R. W. RyR2-dependent modulation of neuronal hyperactivity: A potential therapeutic target for treating Alzheimer’s disease RyR2-dependent modulation of neuronal hyperactivity represents a promising new target for combating AD. J Physiol 602, 1509–1518 (2024).

78. Yuen, S. C., Lee, S. M. Y. & Leung, S. W. Putative Factors Interfering Cell Cycle Re-Entry in Alzheimer’s Disease: An Omics Study with Differential Expression Meta-Analytics and Co-Expression Profiling. Journal of Alzheimer’s Disease 85, 1373–1398 (2022).

79. Yao, J. et al. Increased RyR2 open probability induces neuronal hyperactivity and memory loss with or without Alzheimer’s disease–causing gene mutations. Alzheimer’s and Dementia 18, 2088–2098 (2022).

80. Rattazzi, L. et al. CD4+ but not CD8+ T cells revert the impaired emotional behavior of immunocompromised RAG-1-deficient mice. Transl Psychiatry 3, (2013).

81. Fang, M. et al. Contribution of Rag1 to spatial memory ability in rats. Behavioural Brain Research 236, 200–209 (2013).

82. Qiu, H. & Weng, Q. Screening of Crucial Differentially-Methylated/Expressed Genes for Alzheimer’s Disease. Am J Alzheimers Dis Other Demen 37, (2022).

83. Shen, J.-N.Wang, D.-S. & Wang, R. The Protection of Acetylcholinesterase Inhibitor on β-Amyloid-Induced Injury of Neurite Outgrowth via Regulating Axon Guidance Related Genes Expression in Neuronal Cells. Int J Clin Exp Pathol vol. 5 www.ijcep.com/www.ijcep.com/ (2012).

84. Xiong, M. et al. A γ-adducin cleavage fragment induces neurite deficits and synaptic dysfunction in Alzheimer’s disease. Prog Neurobiol 203, (2021).

85. Urso, K. et al. NFATc3 regulates the transcription of genes involved in T-cell activation and angiogenesis. Blood 118, 795–803 (2011).

86. Ehtewish, H. et al. Profiling the autoantibody repertoire reveals autoantibodies associated with mild cognitive impairment and dementia. Front Neurol 14, (2023).

87. Ka, S. et al. Quantitative proteomics and phosphoproteomics of PPP2R5D variants reveal deregulation of RPS6 phosphorylation through converging signaling cascades. bioRxiv (2023) doi:10.1101/2023.03.27.534397.

88. Tsai, A. P. et al. PLCG2 is associated with the inflammatory response and is induced by amyloid plaques in Alzheimer’s disease. Genome Med 14, (2022).

89. Magno, L. et al. Alzheimer’s disease phospholipase C-gamma-2 (PLCG2) protective variant is a functional hypermorph. Alzheimers Res Ther 11, (2019).

90. Claes, C. et al. The P522R protective variant of PLCG2 promotes the expression of antigen presentation genes by human microglia in an Alzheimer’s disease mouse model. Alzheimer’s and Dementia 18, 1765–1778 (2022).

91. van der Lee, S. J. et al. A nonsynonymous mutation in PLCG2 reduces the risk of Alzheimer’s disease, dementia with Lewy bodies and frontotemporal dementia, and increases the likelihood of longevity. Acta Neuropathol 138, 237–250 (2019).

92. Alam, J. & Scheper, W. Targeting neuronal MAPK14/p38α activity to modulate autophagy in the Alzheimer disease brain. Autophagy vol. 12 2516–2520 Preprint at 10.1080/15548627.2016.1238555 (2016).

93. Yllmaz, Ş.G. et al. Okadaic Acid-Induced Alzheimer’s in Rat Brain: Phytochemical Cucurbitacin E Contributes to Memory Gain by Reducing TAU Protein Accumulation. OMICS 27, 34–44 (2023).

94. Ma, Q., Chen, G., Li, Y., Guo, Z. & Zhang, X. The molecular genetics of PI3K/PTEN/AKT/mTOR pathway in the malformations of cortical development. Genes and Diseases vol. 11 Preprint at 10.1016/j.gendis.2023.04.041 (2024).

95. Gabbouj, S. et al. Altered insulin signaling in Alzheimer’s disease brain-special emphasis on pi3k-akt pathway. Frontiers in Neuroscience vol. 13 Preprint at 10.3389/fnins.2019.00629 (2019).

96. Kitagishi, Y., Nakanishi, A., Ogura, Y. & Matsuda, S. Dietary Regulation of PI3K/AKT/GSK-3β Pathway in Alzheimer’s Disease. http://alzres.com/content/6/3/35.

97. Razani, E. et al. The PI3K/Akt signaling axis in Alzheimer’s disease: a valuable target to stimulate or suppress? Cell Stress and Chaperones vol. 26 871–887 Preprint at 10.1007/s12192-021-01231-3 (2021).

98. Zhang, Y. et al. Vav2 is a novel APP-interacting protein that regulates APP protein level. Sci Rep 12, (2022).

99. Norbury, A. J., Jolly, L. A., Kris, L. P. & Carr, J. M. Vav Proteins in Development of the Brain: A Potential Relationship to the Pathogenesis of Congenital Zika Syndrome? Viruses vol. 14 Preprint at 10.3390/v14020386 (2022).

100. Bai, Y., Xiang, X., Liang, C. & Shi, L. Regulating Rac in the nervous system: Molecular function and disease implication of Rac GEFs and GAPs. BioMed Research International vol. 2015 Preprint at 10.1155/2015/632450 (2015).

101. Ibanez, K. R. et al. Deletion of Abi3/Gngt2 influences age-progressive amyloid β and tau pathologies in distinctive ways. Alzheimers Res Ther 14, (2022).

102. Ghaffari, D., Griffin, J. & St George-Hyslop, P. ABI3 deletion in TgCRND8 mice is associated with reduced amyloid plaque pathology and altered glial response. doi:10.1101/2023.10.05.560956.

103. Bayraktar, A. et al. Revealing the molecular mechanisms of alzheimer’s disease based on network analysis. Int J Mol Sci 22, (2021).

104. Orelio, C. & Dzierzak, E. Expression analysis of the TAB2 protein in adult mouse tissues. Inflammation Research 56, 98–104 (2007).

105. Mosquera-Heredia, M. I. et al. Long Non-Coding RNAs and Alzheimer’s Disease: Towards Personalized Diagnosis. Int J Mol Sci 25, 7641 (2024).

106. Ye, T. et al. Chrysophanol improves memory ability of d-galactose and Aβ25–35 treated rat correlating with inhibiting tau hyperphosphorylation and the CaM–CaMKIV signal pathway in hippocampus. 3 Biotech 10, (2020).

107. Ye, S. et al. Genistein protects hippocampal neurons against injury by regulating calcium/calmodulin dependent protein kinase IV protein levels in alzheimer’s disease model rats. Neural Regen Res 12, 1479–1484 (2017).

108. Müller, M., Cárdenas, C., Mei, L., Cheung, K. H. & Foskett, J. K. Constitutive cAMP response element binding protein (CREB) activation by Alzheimer’s disease presenilin-driven inositol trisphosphate receptor (InsP 3R) Ca 2+ signaling. Proc Natl Acad Sci U S A 108, 13293–13298 (2011).

109. Park, D. et al. Activation of CaMKIV by Soluble Amyloid-b 1-42 Impedes Trafficking of Axonal Vesicles and Impairs Activity-Dependent Synaptogenesis. https://www.science.org (2017).

110. Kamboh, M. I. et al. Genome-wide association study of Alzheimer’s disease. Transl Psychiatry 2, (2012).

111. Ramanan, V. K. et al. Variants in PPP2R2B and IGF2BP3 are associated with higher tau deposition. Brain Commun 2, (2020).

112. Zhou, Z. et al. Downregulation of PIK3CB Involved in Alzheimer’s Disease via Apoptosis, Axon Guidance, and FoxO Signaling Pathway. Oxid Med Cell Longev 2022, (2022).

113. Deng, L., Zhang, J., Cao, K., Shang, M. & Han, F. Combining GEO Database and the Method of Network Pharmacology to Explore the Molecular Mechanism of Epimedium in the Treatment of Alzheimer’s Disease. in ACM International Conference Proceeding Series 522–530 (Association for Computing Machinery, 2022). doi:10.1145/3581807.3581884.

114. Junyent, F. et al. Gene expression profile in JNK3 null mice: A novel specific activation of the PI3K/AKT pathway. J Neurochem 117, 244–252 (2011).

115. Wegrzyn, D., Zokol, J. & Faissner, A. Vav3-Deficient Astrocytes Enhance the Dendritic Development of Hippocampal Neurons in an Indirect Co-culture System. Front Cell Neurosci 15, (2022).

116. Nahalka, J. 1-L Transcription in Alzheimer’s Disease. Curr Issues Mol Biol 44, 3533–3551 (2022).

117. Bonham, L. W. et al. Neurotransmitter Pathway Genes in Cognitive Decline During Aging: Evidence for GNG4 and KCNQ2 Genes. Am J Alzheimers Dis Other Demen 33, 153–165 (2018).

118. Bayraktar, A. et al. Revealing the molecular mechanisms of alzheimer’s disease based on network analysis. Int J Mol Sci 22, (2021).

